# Evaluating assembly and variant calling software for strain-resolved analysis of large DNA-viruses

**DOI:** 10.1101/2020.05.14.095265

**Authors:** Z.-L. Deng, A. Dhingra, A. Fritz, J. Götting, P. C. Münch, L. Steinbrück, T. F. Schulz, T. Ganzenmüller, A. C. McHardy

## Abstract

Infection with human cytomegalovirus (HCMV) can cause severe complications in immunocompromised individuals and congenitally infected children. Characterizing heterogeneous viral populations and their evolution by high-throughput sequencing of clinical specimens requires the accurate assembly of individual strains or sequence variants and suitable variant calling methods. However, the performance of most methods has not been assessed for populations composed of low divergent viral strains with large genomes, such as HCMV. In an extensive benchmarking study, we evaluated 15 assemblers and six variant callers on ten lab-generated benchmark data sets created with two different library preparation protocols, to identify best practices and challenges for analyzing such data.

Most assemblers, especially metaSPAdes and IVA, performed well across a range of metrics in recovering abundant strains. However, only one, Savage, recovered low abundant strains and in a highly fragmented manner. Two variant callers, LoFreq and VarScan2, excelled across all strain abundances. Both shared a large fraction of false positive (FP) variant calls, which were strongly enriched in T to G changes in a “G.G” context. The magnitude of this context-dependent systematic error is linked to the experimental protocol. We provide all benchmarking data, results and the entire benchmarking workflow named QuasiModo, **Quasi**species **M**etric **d**etermination **o**n **o**mics, under the GNU General Public License v3.0 (https://github.com/hzi-bifo/Quasimodo), to enable full reproducibility and further benchmarking on these and other data.

## Introduction

Human cytomegalovirus (HCMV) causes a lifelong infection that is typically without major clinical symptoms. After primary infection HCMV persists latently in infected cells [1]. Primary or (re-)infections and reactivation of HCMV can cause significant morbidity and severe complications in immunocompromised individuals, such as HIV-infected persons, transplant recipients or congenitally infected children [2,3]. HCMV has a double-stranded DNA genome of approximately 235 kb, including terminal and internal repeats, which contains at least 170 open reading frames [4]. With genome sizes of known viruses ranging from ∼1 kb (*Circovirus SFBeef*) to 2 Mb (*Pandoravirus salinus*) [5], HCMV belongs to the larger known viruses and has co-evolved with its host for millions of years [6]. Multiple HCMV strain infections (i.e. with more than one strain at the same time) probably contribute to prolonged viremia, delayed viral clearance and other complications [7–10].

The establishment of high-throughput sequencing techniques and accompanying bioinformatics analysis methods has greatly advanced viral genomic research [11–16]. Assembling viral genomes of individual virus strains from a mixed population and variant calling are essential for characterizing the evolution and genetic diversity of viral pathogens such as HCMV *in vivo*. Although HCMV mutates and evolves more slowly than many RNA viruses and not any faster than other herpes viruses, high levels of genetic variation due to mixed (i.e. multiple) viral strain infections in an individual are often observed [17–20]. These multiple strain infections likely result from reactivation of latent strains and/or re-infections [17,21,22].

Assemblers leverage short read sequence data by linking sequences using kmer or read graphs, and, in some cases, variant frequencies, to reconstruct viral haplotypes, such as the recently developed HaROLD [23], which makes use of longitudinal sequence data. There are also many variant callers available, including programs for calling low-frequency variants, such as LoFreq [24], VarScan2 [25], and the commercial CLC Genomics Workbench [26]. Those programs use information on basecall and mapping quality to determine if a variant site in a read may be due to sequencing error, mapping bias or reflects true biological diversity [24–26].

A recent study on simulated and mock viromes suggests that the choice of assembler largely influences virome characterization [27]. Several assemblers that we evaluated, including IDBA-UD [28], SPAdes [29], Ray [30] and Megahit [31], were previously assessed on more divergent, simulated and spiked mock viromes [27,32]. This in one case included strains of less than 97% average nucleotide identity (ANI) [33], which resulted in shorter assemblies for low divergent community members. Viral haplotype assemblers reconstruct small viral genomes, such as HIV, Zika and hepatitis C virus, with good genome fractions. However, these may be highly fragmented, in case of Savage [34], or consist of longer contigs with a substantial amount of misassemblies, in case of PEhaplo [35], QuasiRecomb [36] and PredictHaplo [37]. Viral haplotype assemblers have so far been mostly evaluated on much smaller and more divergent genomes (genome size around 10 kb with divergence of up to 12.7%) [34,38]. They have not been assessed on substantially larger genomes with low density of variants so far. A recent assessment of variant callers [39] reported variable, in part complementary performances of FreeBayes [40], LoFreq, VarDict [41], and VarScan2 in minority variant detection on simulated short read data from Respiratory Syncytial Virus (RSV), which is a small virus with a 15 kb genome size.

So far, strain-level assembly and variant calling methods have not been evaluated for large DNA viruses, where runtime and memory consumption of the algorithms might also be critical, nor on benchmark data that include experimental biases of library preparation and sequencing. To investigate these issues, in the largest benchmark of its kind so far, we created and sequenced ten samples of HCMV strains with different mixing ratios and then evaluated 21 computational methods on the resulting WGS data. Analysis of these lab-created benchmark data sets allowed us to dissect the effects of computational methods and library preparation protocols.

## Results

### Creation and quality control of viral sequence samples

To produce a benchmark dataset of mixed viral strains that also includes technical artifacts introduced in experimental data generation, we created viral strain mixtures mimicking clinical samples from patients with mixed strain infections *in vitro*. For this, we combined viral DNA of the HCMV strains TB40/E BAC4 and AD169 (designated as “TA”), derived directly from bacterial artificial chromosomes (BAC) with these viral genomes and prepared from *Escherichia coli*, or the strains TB40/E BAC4 and Merlin (designated as “TM”), which were amplified in human cell-cultures, respectively, at mixing ratios of 1:1, 1:10 and 1:50. The ANIs between each pair of those strains are around 0.977 (Table S1). In addition, pure strains were sequenced separately in each experiment, resulting in four data sets with the TB40/E and AD169 strains without target enrichment and the TB40/E and Merlin strain after enrichment. For the TA mixture experiments, we used a library preparation protocol (protocol 1, details in Material and Methods) without target enrichment, for the TM mixtures a protocol including target enrichment (protocol 2). All ten samples (6 HCMV strain mixtures and 4 pure strains) were sequenced using 2x 300 bp paired-end sequencing (Illumina MiSeq), resulting in 1.58 million raw reads on average per sample. After quality control, 1.1 million quality reads per sample with average base quality above 30 remained.

As the HCMV strains for the TA mixtures and corresponding pure strain samples were extracted from *E. coli* BACs, *E. coli* reads were found in those samples with an average fraction of 48.6±16.5% (Table S2). Based on the genome size of HCMV (235K) and *E. coli* (4.6M), the abundance of contaminating *E. coli* is thus around 5%. The three TM data sets and the pure Merlin strain, TM-0-1, did not include detectable bacterial contamination, but 51.7% of the reads of TM-1-0 (pure TB40/E strain) were of human origin.

### Strain-resolved genome assembly

For mixed strain data sets, the ultimate aim for assembly is to recover the genomes of individual strains. To obtain a comprehensive performance overview for existing software, we evaluated the performances of the generic (meta-)genome assemblers SPAdes, metaSPAdes [42], Megahit, ABySS [43], Ray, IDBA-UD, Tadpole, which is a part of the BBMAP toolkit [44] and IVA [45], Vicuna [46], as well as the viral haplotype assemblers Savage, PredictHaplo, PEhaplo, QuasiRecomb, ShoRAH [47] and VirGenA [48] on our data sets (Material and Methods).

Assemblies were assessed based on common assembly quality metrics with metaQUAST [49], such as genome fraction, duplication ratio, largest alignment, NGA50 using both strain genomes as references for the respective mixtures (Methods, Figure 1). The genome fraction is defined as the fraction of the reference genome covered by at least one contig. The duplication ratio is the number of bases of the reference genome covered, divided by the total number of aligned bases from the assembly. The largest alignment is the size of the biggest contig that aligned to the reference genome. The NGA50 value of an assembly is calculated by first sorting the aligned contigs, after being split at misassembly events, by size in descending order and returning the length of the contig that exceeds 50% genome fraction. If an assembler fails to produce 50% genome fraction, the NGA50 value cannot be calculated and was set to 0 kb. To further summarize the performance of assemblers on the HCMV datasets, we defined a composite quality metric for strain-resolved assembly performances, consisting of a weighted score combining the metaQUAST assembly metrics “duplication ratio”, “genome fraction”, “largest alignment”, “NGA50”, “number of contigs”, and “number of mismatches per 100 kb” (Materials and Methods). In this weighted score, we considered genome fraction and largest alignment the most important metrics, since they reflect the ability of the assembler to reconstruct individual strains and the completeness of the largest assembly.

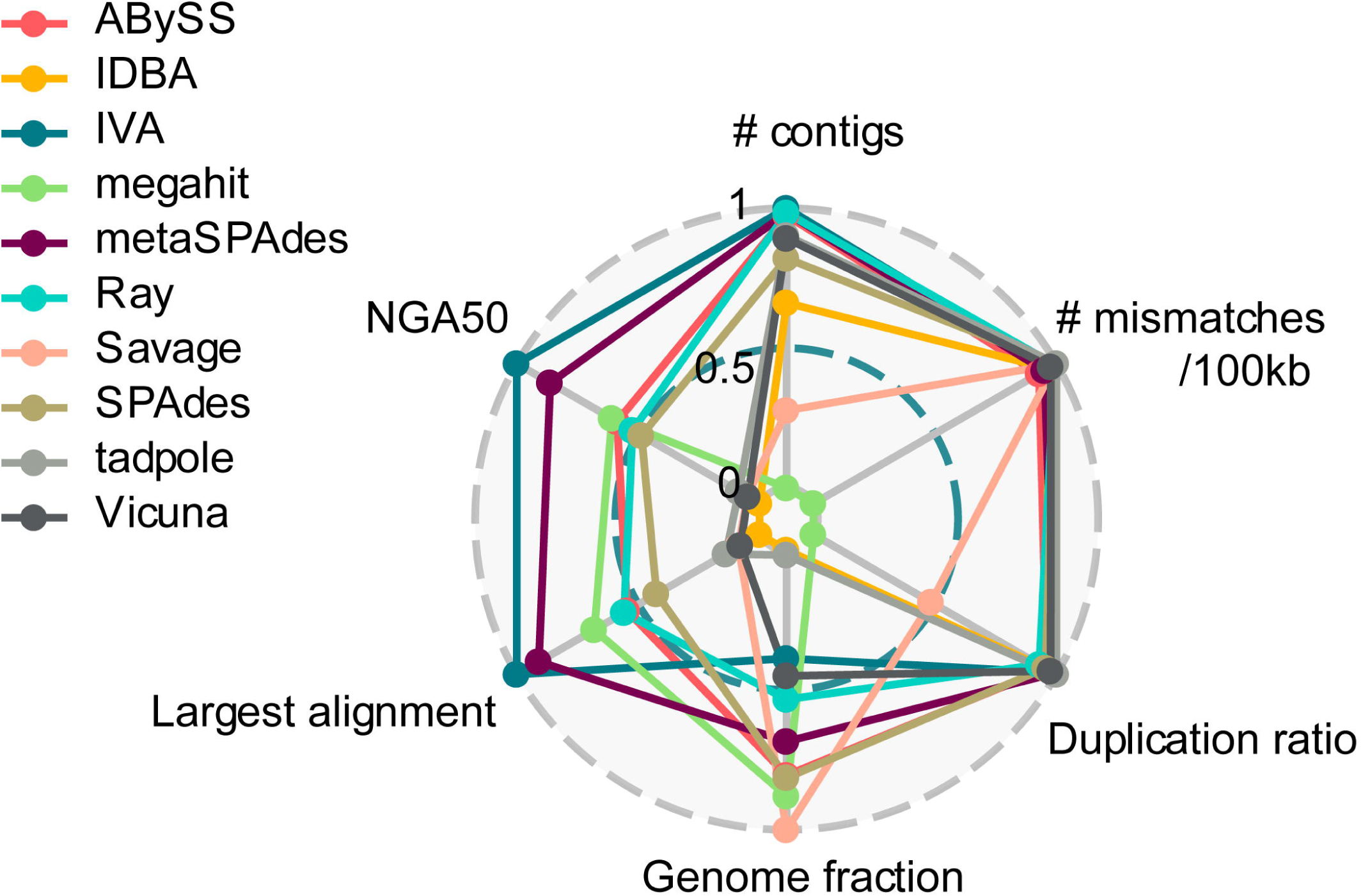

All programs reconstructed the genome sequence much better for the dominant than for the minor strain. With the weighted summary score, metaSPAdes achieved the highest score (8.57), with a large genome fraction assembled (54.5±6.4% versus 45.3±12.4% for IVA, mean ± standard deviation) and second best for largest alignment (145.9±56.6 kb), NGA50 (102.3±69.0 kb), number of contigs (12.5±9.0) and duplication ratio (1.01±0.01) (Figure 1, Table S3-4). Next were IVA (8.12), which was ranked best for largest alignment, NGA50, and number of contigs, and ABySS (7.50) (Figure 1, Table S3-4). IVA produced on average the fewest (8.1±7.9), and longest contigs (159.6±77.8 kb), especially for abundant strains (160.8±72.3 kb) (1/0, 50/1, 10/1), with only very few parts of the genomes covered multiple times (duplication ratio of 1.01±0.03) (Figure 1-2, Table S3). The Tadpole assembly had the lowest duplication ratio (1.001±0.001) and the fewest mismatches per 100 kb (32.2±54.4, Figure 1-2, Table S3). However, this was mainly because it assembled very little data and generated short contigs (NGA50 10.9±15.4 kb) that covered less than half (33.8±15.6 %) of the underlying genomes.

The haplotype assembler Savage in reference-based mode recovered the most (64.4±27.2% genome fraction) of both strains, even for the low abundant ones (1/10, 1/50) (Figure 2). However, it produced shorter contigs (largest contig length 21.5±22.8 kb) and many duplicates (1.38±0.23). Megahit recovered most (93.4±5.7%) of the genome sequence for the dominant strains, followed by ABySS (92.7±3.1%), SPAdes (91.8±4.1%) and Ray (91.6±7.8%), however much less for the low abundant strains (38.3±20.0%, 37.3±14.5%, 35.2±6.6%, 12.2±4.5%, respectively). MetaSPAdes and IVA also recovered a relatively large fraction (83.4±28.2% and 86.4±21.2%, respectively) of the dominant strains, but only little (38.7±28.4% and 8.5±6.0% respectively) of the genome for low abundant strains.

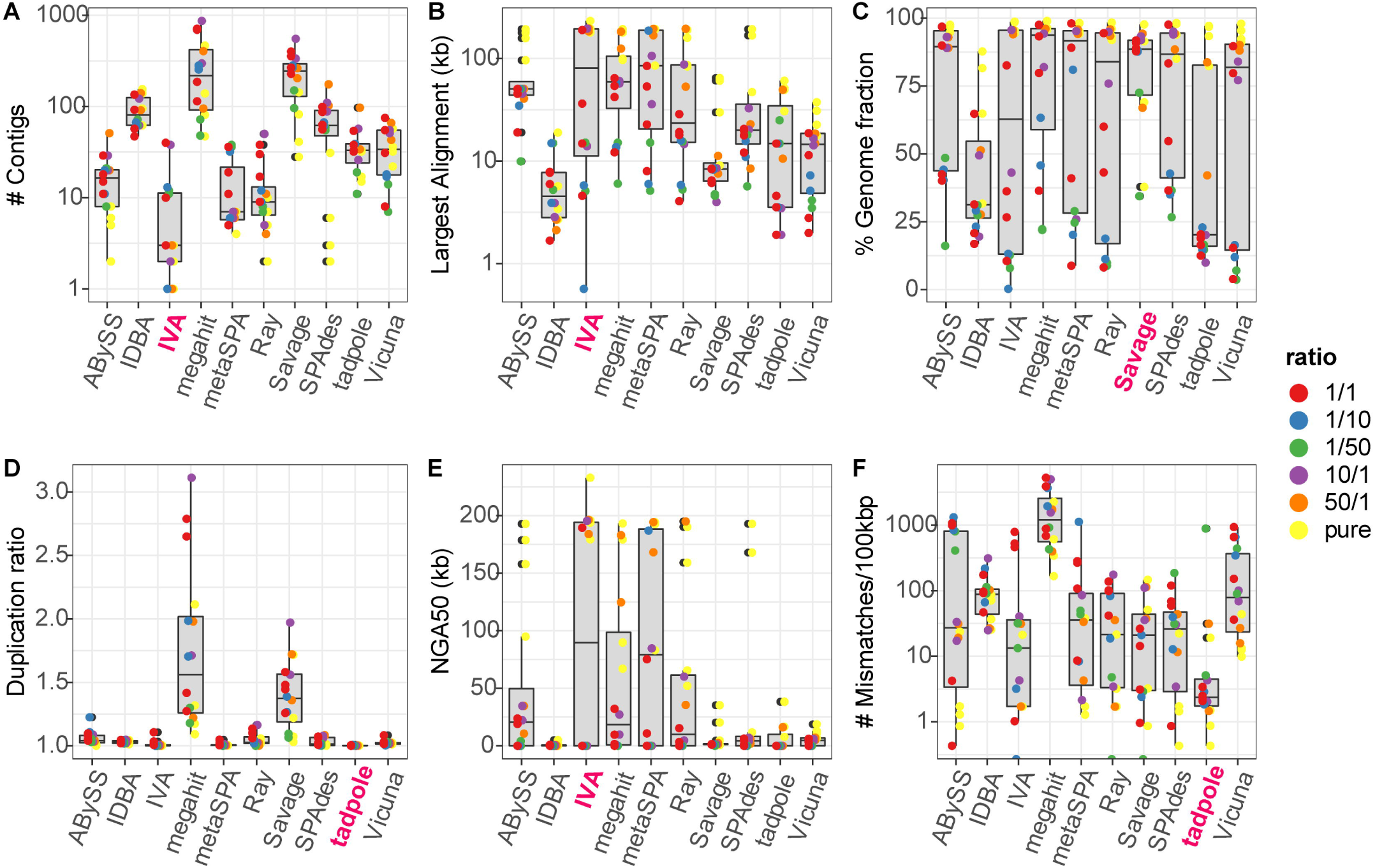

All other haplotype assemblers, *i.e.* Savage in *de novo* mode, PredictHaplo, PEhaplo, QuasiRecomb, ShoRAH and VirGenA, assembled no contigs and were terminated after running for more than 10 days using 24 CPU cores. Furthermore, we also tested 1000 random weights sets to calculate the summary score, and the top two assemblers (metaSPAdes and IVA) maintained this ranking for ∼850 out of 1000 sets. This suggests that the assemblers with good performance deliver a high quality assembly across most metrics.

As genome assembly can be computationally intensive and time consuming, we also benchmarked the disk space consumption (IO output), memory (maximum memory requirement) and run time of the different algorithms. Ray and ABySS used less than 300 MB for the output while IVA, SPAdes, metaSPAdes and Savage consumed more than 20 GB of disk space for output or intermediate output (Figure 3). Megahit was the most memory efficient assembler, using less than 1 GB memory, whereas ABySS, Savage and Vicuna consumed more than 10 GB. As to the run times, Megahit required around ten minutes for each assembly, while Vicuna and Savage needed more than 20 hours on a server with sixty-two 2.4GHZ CPUs, 200 TB disk space and 1 TB memory.

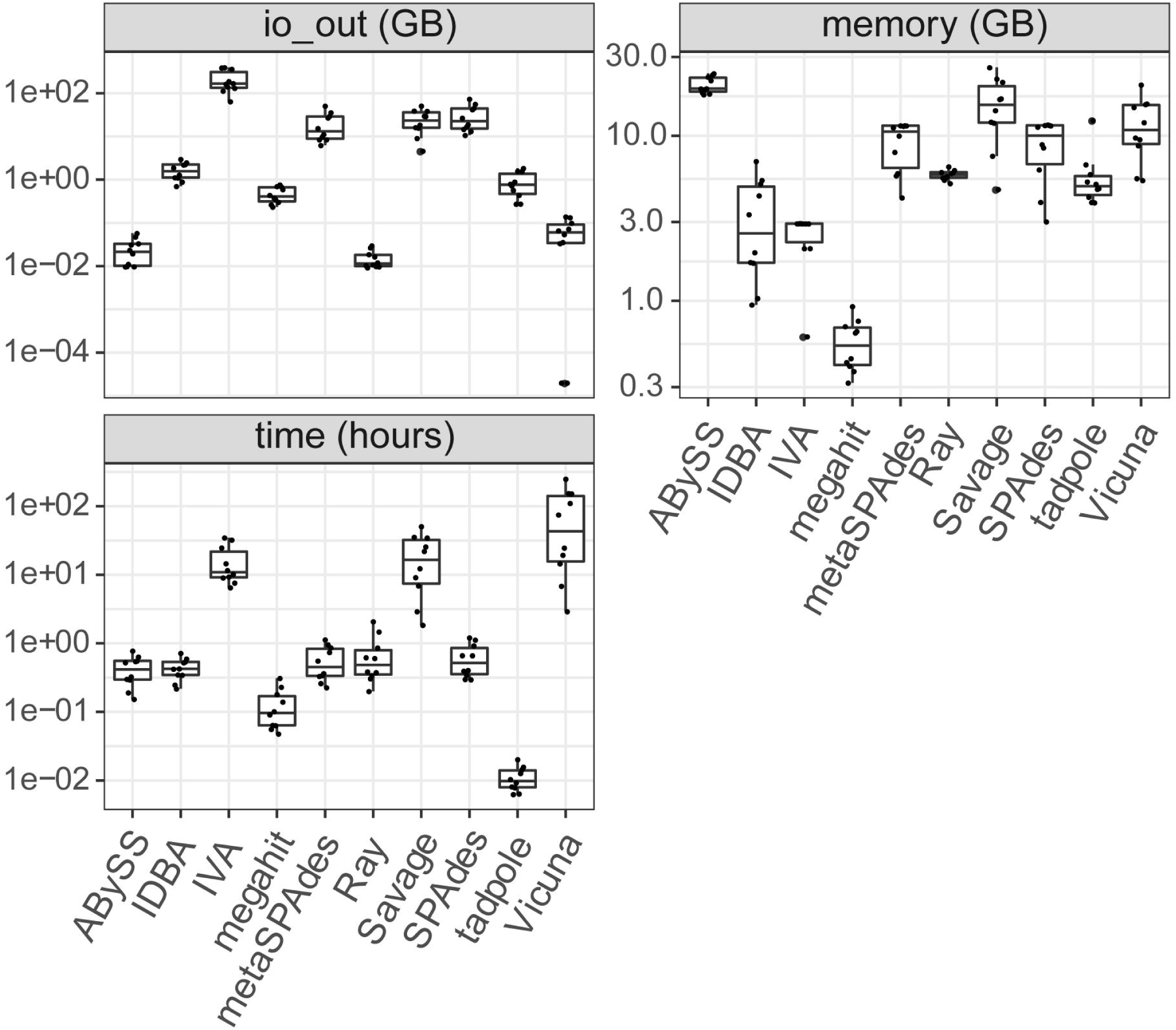

### Variant calling

We evaluated the variant callers LoFreq, VarScan2, the low frequency variant caller of the CLC genomics workbench, BCFtools [50], FreeBayes and the GATK HaplotypeCaller [51] on the six mixed strain and four pure strain (three different strains, details in material and methods) WGS samples. A ground truth was generated by pairwise genome alignment of the respective strains with MUMmer [52] (Methods, Figure 4), which identified around 3500-4000 variants, including ∼200 short insertions and deletions (InDels). Sites in these genomes were then classified as variant or non-variant in this alignment, and compared to predicted variants, to determine true positive (TP), false negative (FN) and false positive (FP) calls. Since the major strain in each mixture was used as reference, we could evaluate the performance of those variant callers in identifying low frequency variants originating from the minor strain in the mixture, with the expected low frequency variants being 2% and 10%, respectively, in the mixtures with ratios of 1:50 and 1:10. Variant calls for which a false nucleotide was predicted for a variant site were also considered as false positives. Based on the number of TP, FN and FPs we calculated precision, recall and the F1-score as detection quality metrics for each caller and sample. Precision, or purity, reflects the fraction of predicted variants that are true variants: 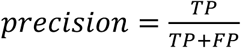; it thus quantifies how reliable the predictions of a particular method are. Recall is sometimes also known as completeness, and measures the fraction of truly existing variants in a data set that have been detected by a caller 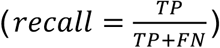, it thus measures how complete the predictions of a caller are with respect to the variants that are there to discover. To allow a comparison based on a single metric, the F1-score is commonly used, which is the harmonic mean of precision and recall, i.e. 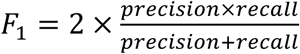.

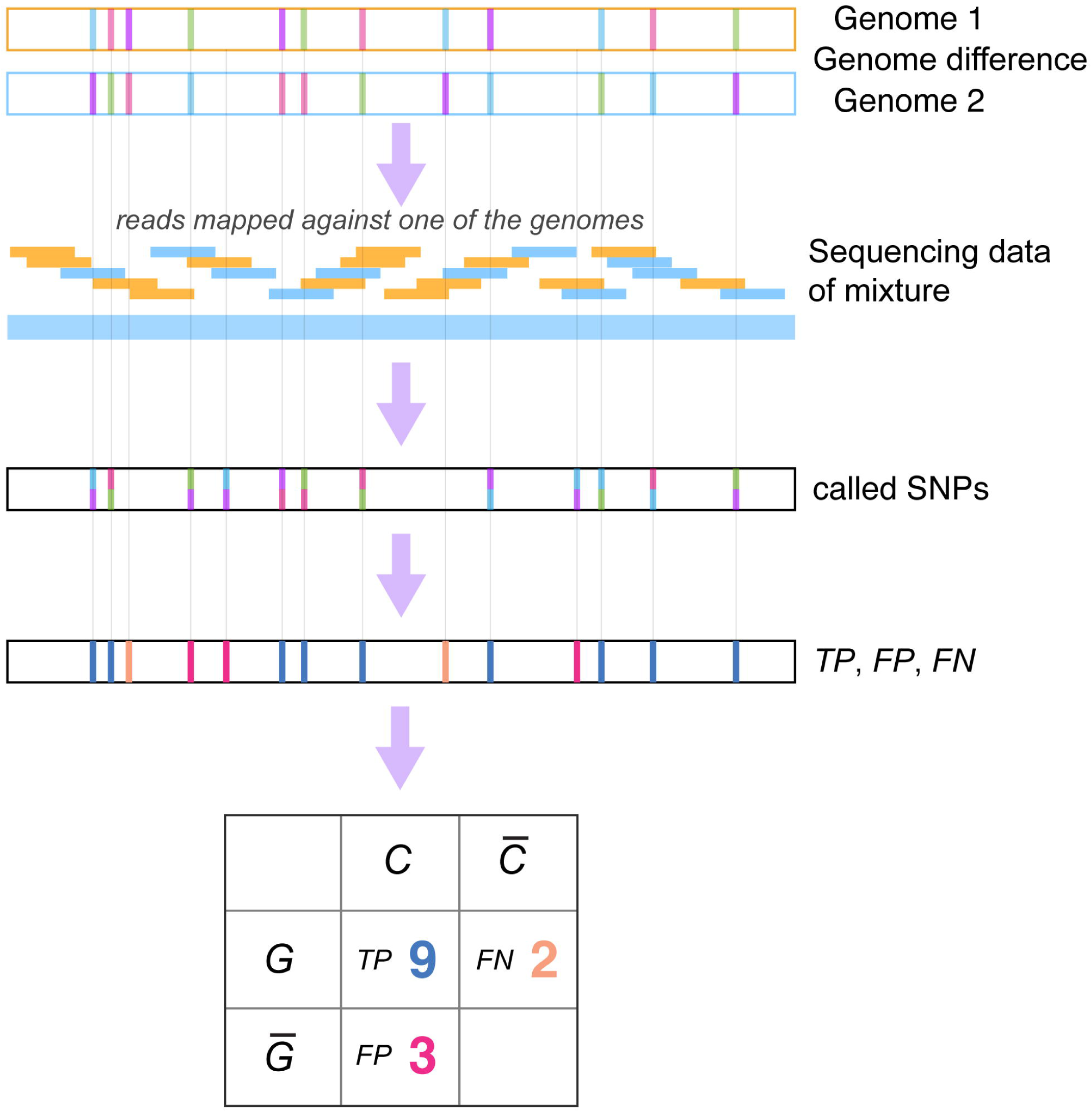

Applying the commonly used cutoff of 20 for Phred quality scores (QUAL) [53] for accepting predicted variants, we evaluated the performance of variant callers on single nucleotide polymorphisms (SNPs). LoFreq achieved the best average precision (0.940±0.011) and VarScan2 the highest recall (0.872±0.050, Figure 5A, Table S5) across mixture samples. LoFreq and VarScan2 consistently performed best across samples, with average F1-scores, of 0.890±0.009 and 0.880±0.011, respectively (Figure 5B, Table S5). CLC had a slightly lower F1-score (0.806±0.025), and was more variable in performance across samples, while BCFtools, GATK and Freebayes performed poorly (F1-score: 0.166±0.288, 0.261±0.388 and 0.289±0.428, respectively), particularly due to low recall (0.122±0.230, 0.215±0.338 and 0.253±0.386). Across all strains and abundance ratios tested, LoFreq consistently performed well, while VarScan2 was consistent across abundance ratios but performed differently for the two strain mixtures (varied in precision) and CLC’s recall dropped dramatically for mixture TM-1-50. BCFtools, GATK and FreeBayes performed poorly in comparison for all samples, and for highly diluted samples, their recall was almost 0. To analyze the effect of their returned Phred quality scores on variant callers’ performances, we evaluated both SNPs and InDels called with different thresholds for their quality scores using a recall-precision curve. LoFreq had the best recall-precision balance followed by VarScan2 and CLC, while FreeBayes demonstrated high performance on samples TA-1-1 and TM-1-1 (Figure S1). To compare variant caller performances under optimized performance conditions, we also determined performances of variants called using the best F1-scores over these different settings across all samples. Notably, the performance of FreeBayes increased substantially, and that of CLC slightly, while the performances of other methods remained similar (Figure 5C and 5D, Figure S2, Table S6).

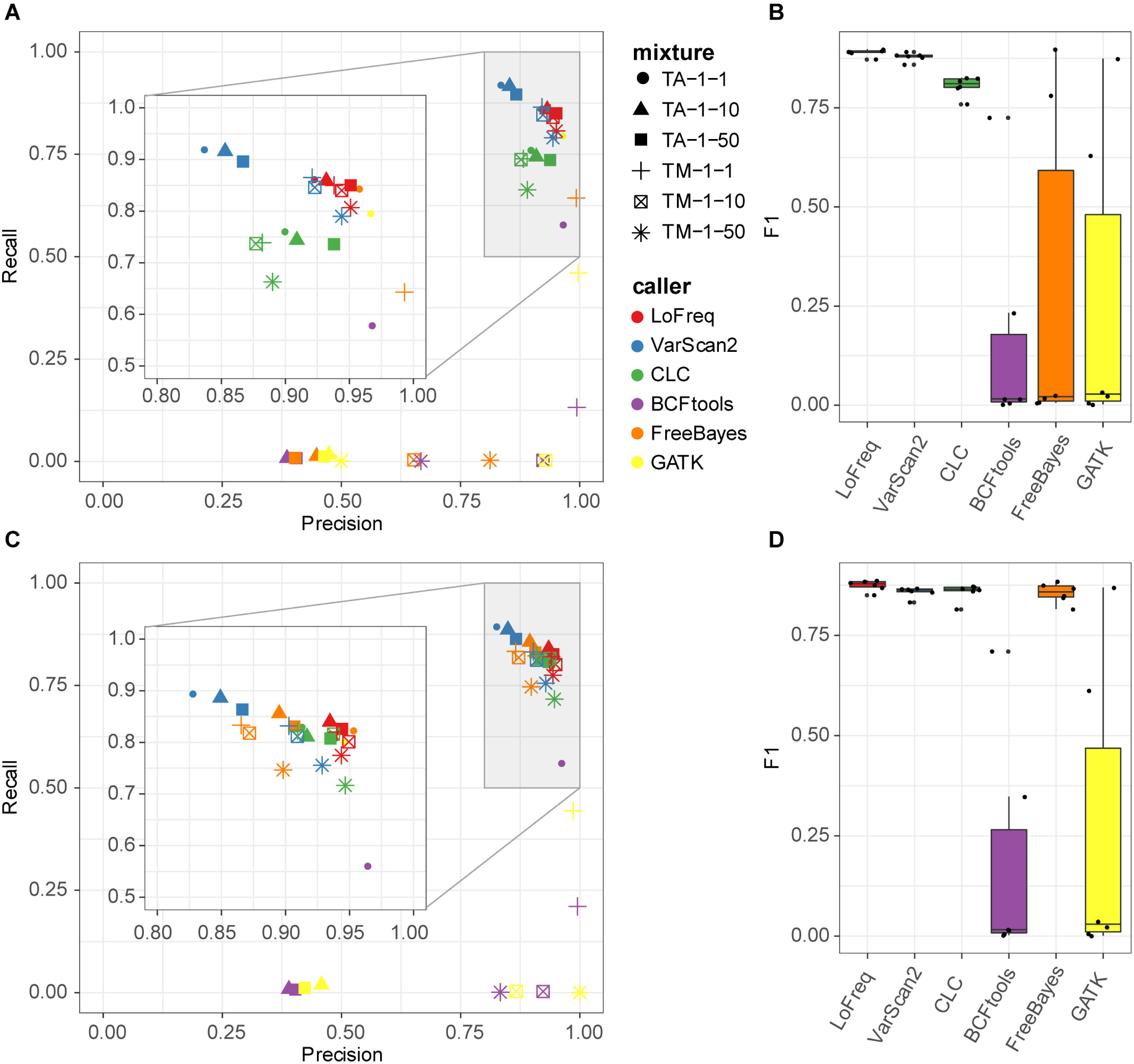

The callers achieving good recall, LoFreq, CLC and VarScan2, identified around 2400 to 2700 shared true positive SNPs from all mixed strain samples when using a quality score threshold of 20 (Figure 6). On the pure strain samples, where no SNPs were expected, LoFreq and VarScan2 predicted 61±33 and 71±42 false positives, respectively, substantially less than for the mixed strain samples (164±59 and 381±163). Notably, of these false positives in mixtures, 70.7±17.3% (based on LoFreq predictions) and 37.6±7.9% (based on VarScan2) were shared (Figure S3). This significant overlap (Fisher’s exact test p-value <2.2×10-16, odds ratio 3416.8±1601.1), indicates a systematic shared bias regardless of variant callers. Variant calling (Figure S4) indicated that allele frequencies intended by dilutions were closely reached with protocol 2 (TM mixture) and differed slightly more for protocol 1 (TA mixture).

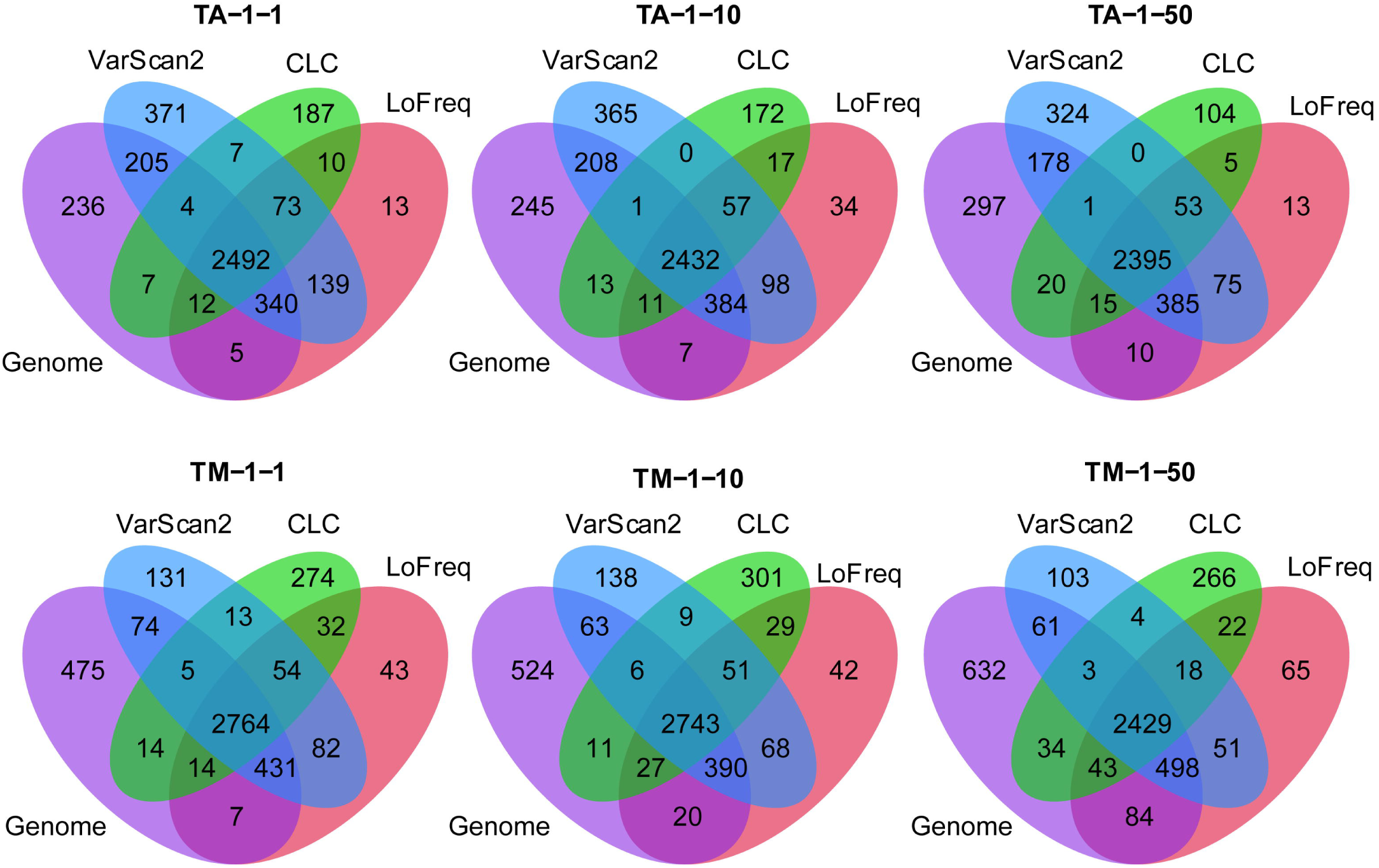

### Genomic context of variant calls

We analyzed whether there was a specific genomic signal associated with variant calls, considering separately correct and false calls using mutational context analysis [54–56]. Focusing exemplarily on LoFreq, this approach analyzes the frequency of a certain SNP together with its sequence context, specifically the flanking3’ and 5’ bases. For the predictions of a certain caller, the genomic context of the six substitution types (C to A, C to G, C to T, T to A, T to C and T to G) was calculated with the R package SomaticSignatures [56] for the six mixtures and four pure strain samples (2 samples of TB40/E, 1 of Merlin, and 1 of AD169). Since the analysis is not strand-specific, the above were considered equivalent with G to T, G to C, G to A, A to T, A to G and A to C, respectively. We observed a strong, context-independent preference for C to T or T to C transitions (with a fraction of 0.803±0.016 of all variant calls across samples; top panel of Figure 7A and Figure 7B), which was even more pronounced for the true positives (middle panel of Figure 7A and Figure 7B), but not for FPs. For variants observed across pairwise combinations of 30 *E. coli* and 30 HIV genomes, which were obtained from NCBI RefSeq database (Table S7), respectively, we observed concordant results (Figure S5-S6). For these data, transitions accounted for 0.716±0.058 and 0.681±0.017 of variants between genome pairs, respectively.

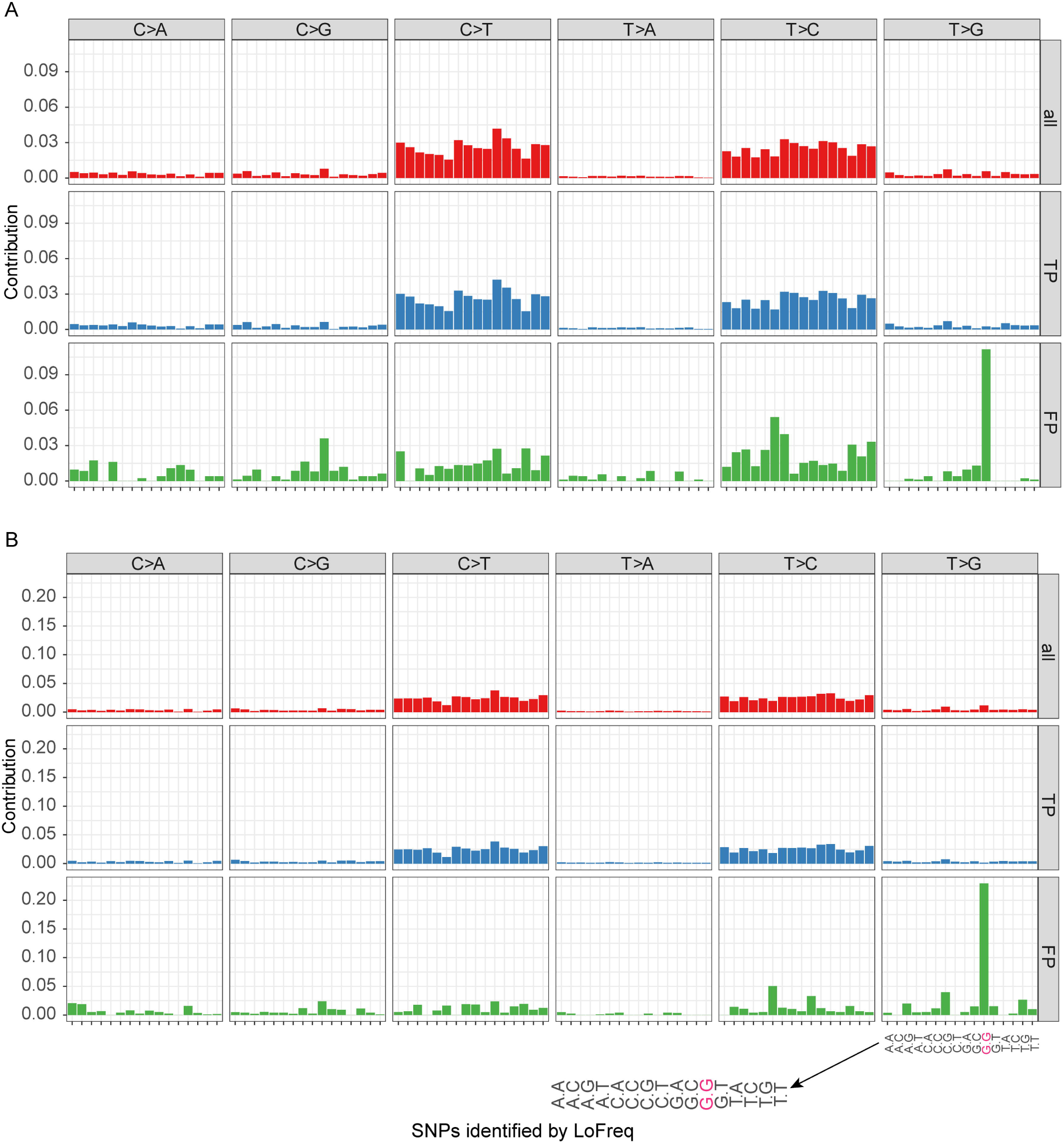

We found a pronounced context dependent signal for false positive calls of LoFreq and VarScan2. Here, T to G variants in a G.G context correspond mostly to FPs in the TA and TM mixtures (57.1±10.0% and 86.8±18.3%, respectively; bottom panel of Figure 7A and Figure 7B). This enrichment is highly significant (p-value <0.0001, Fisher’s exact test), with an odds ratio of around 45.2 for the TM mixture; i.e. T to G calls are 45.2 times more frequent in this context than in others and 19.9 more frequent for the TA mixture. For false variant calls on the pure Merlin and AD169 samples, T to G calls in a G.G context were even more dominant. For LoFreq on the pure Merlin (TM-0-1) sample, the genomic context pattern of false calls is highly correlated with the context pattern of false positives for all mixed strain samples, with an average Pearson correlation of 0.903 (p-value <0.0001). For the AD169 strain and respective mixtures, this correlation (Pearson) is lower, on average 0.697, but still highly significant (p-value <0.0001).

The allele frequencies of the FP LoFreq variants were substantially lower than those of the true positive variants (Figure S7, Wilcoxon test p-value <2.2×10-16), except for the TA-1-50 sample, which had the highest-level *E. coli* cloning vector contamination. False T to G calls in a G.G context had a lower frequency than other false calls (p-value 1.181×10-10 for TM mixtures: Figure S8, 8.16×10-10 for TA mixtures). The allele frequency of those FP SNPs was slightly lower in protocol 2 (TM, 0.0237±0.0522) than in protocol 1 (TA, 0.0242±0.0121) with a Wilcoxon p-value = 0.000559, 95% CI = [0.00363, 0.0120]. The extent of the signal differed between samples created with different protocols. Though the overall FP rate was similar, the context-dependent false calls T to G in G.G doubled in protocol 2 (Figure 7A, 7B). We found no such signal for false LoFreq variants calls on MiSeq sequencing data from HIV lab data [57], even though the frequency of GTG/CAC patterns in both genomes are similar (Figure S9).

## Materials and methods

### Creation and sequencing of HCMV strain mixtures

We created mixtures for two pairs of strains: “TB40/E BAC4” with “AD169 subclone HB5” (TA) and TB40/E BAC4 with strain Merlin (TM). For each strain pair, mixtures with three different mixing ratios, 1:1, 1:10 and 1:50, were created. Accordingly, strains “AD169” and “Merlin” are the dominant strains in the mixtures, and their genomes were used as reference for variant calling in mixed samples. In addition, the pure strains were sequenced. The name of the mixture specifies the included strains and the mixing ratio. For instance, a mixture of TB40/E and Merlin with a ratio of 1:10 is denoted by TM-1-10. Pure strain samples are denoted as TA-1-0 for TB40/E and TA-0-1 for AD169, which were created with protocol 1 (details, see below), as well as TM-1-0 for TB40/E and TM-0-1 for Merlin, created with protocol 2.

Two protocols were used to generate the sequencing libraries. In protocol 1, the DNA of TA mixtures (TA-1-1, TA-1-10 and TA-1-50) and pure strain samples (TA-0-1, TA-1-0) was extracted from the BAC host *E. coli* strain GS1783 using the Plasmid Midi Kit (Macherey Nagel). Library preparation was performed using an Ultra II FS-Kit from NEB according to the standard protocol from the manufacturer. Fragmentation time was 10 minutes and the library was amplified 4 cycles for the mixtures and 5 cycles for the pure BACs, multiplexed and sequenced on a MiSeq (Illumina) using reagent kit v3 to generate 2 × 300 bp paired-end reads.

Protocol 2 was used to generate the TM mixture data sets (TM-1-1, TM-1-10, TM-1-50) and the pure strain samples data sets (Merlin, TM-0-1 and TB40/E BAC4, TM-1-0). The HCMV strains TB40/E BAC4 and Merlin were isolated from cell cultures. The library preparation was performed as we previously described [20] with the KAPA library preparation kit (KAPA Biosystems, USA) with a few modifications. After PCR pre-amplification (6-14 cycles) with adapter specific primers, up to 750 ng of DNA was target enriched for HCMV fragments using HCMV specific RNA baits. HCMV enriched libraries were indexed, amplified (17 to 20 cycles) using TruGrade oligonucleotides (Integrated DNA Technologies), multiplexed and sequenced on a MiSeq (Illumina) using reagent kit v3 to generate 2 × 300 bp paired-end reads.

### Quality control of the sequencing data

Sequencing reads produced by the MiSeq sequencer were quality controlled using fastp v0.19.4 [58]. Fastp is an all in one FASTQ data preprocessing toolkit with functionalities including quality control, adapter detection, trimming, error correction, sequence filtering and splitting. The remaining adapter sequences were clipped from the raw reads as well as bases at the 5’ or 3’ of the reads with a base quality score of less than 20. Reads shorter than 130 bp after trimming were removed. The remaining PhiX sequences (originating from the Illumina PhiX spike-in control) were also removed from the dataset by mapping all quality-controlled reads against the PhiX reference genome downloaded from Illumina using BWA-MEM v0.7.17 [59]. Contamination from *E. coli* and the human host were also removed using the same method.

### Consensus assembly and evaluation

To benchmark the performance of commonly used assemblers, we evaluated SPAdes v3.12.0 (with kmer sizes: 21, 33, 55, 77, 99, 127 and --careful option), metaSPAdes v3.12.0 (kmer sizes: 21, 33, 55, 77, 99, 127), Megahit v1.1.3 (kmer sizes: 21, 41, 61, 81, 101, 121, 141, 151,), Ray (kmer size: 31), ABySS v2.1.4 (kmer size: 96), IDBA v1.1.3 (default settings), Tadpole v37.99 (default settings), IVA v1.0.9 (default settings) and Vicuna v1.3 (default settings). The quality of the resulting contigs or scaffolds was then assessed with metaQUAST v5.0.2. Only contigs longer than 500 bp were taken into account. Since the reference genomes of those strains are highly similar, with an ANI around 98%, only unique mappings were considered in the assessment, i.e. not allowing a single contig to map to both reference genomes in the combined reference report. The metrics include the overall number of aligned contigs, the largest alignment, genome fraction, duplicate ratio, NGA50, number of mismatches per 100 kb. Here, “largest alignment” refers to the largest contig or scaffold that mapped to the reference genome. “Genome fraction” represents the fraction of the genome recovered by contigs from an assembly. The “duplication ratio” is the total number of aligned bases in the assembly divided by the total number of those in the reference (https://github.com/ablab/quast). NGA50 is the N50 value of the contigs that mapped to the reference genomes with contigs being split at misassemblies. The NGA50 value cannot be calculated for the assemblies which recover less than 50% of the genome in terms of genome fraction and was set to 0 instead to ensure comparability. The individual reference report from metaQUAST was used to evaluate the performance for abundant or low abundant strains in mixtures. All overall metrics values regardless of the specific strain in the mixture were calculated using the combined reference report from metaQUAST, except for NGA50.

### Haplotype reconstruction

Of viral quasispecies assemblers, we ran PEHaplo v0.1, PredictHaplo v0.4, Savage v0.4.0, QuasiRecomb v1.2, ShoRAH v1.9.95 and VirGenA v1.4 using default settings (for details see the code repository). We did not run HaROLD, as this requires longitudinal clinical samples from the same source. The haplotype assemblies were evaluated using metaQUAST together with the consensus assemblies mentioned above.

### A composite quality metric for strain-resolved assembly

To summarize assembly performances, we defined a weighted score based on the metaQUAST assembly metrics using combined reference including genome fraction, largest alignment, NGA50, duplication ratio, number of contigs, and number of mismatches per 100 kb. As NGA50 is not available in the combined reference report of metaQUAST, we used the average NGA50 based on individual genomes from the individual references report. Of these metrics, we considered genome fraction and largest alignment as the most important metrics, since they reflect the ability of the assembler to reconstruct individual strains. To calculate a weighted summary score for assembler performance, we weighted the above metrics by the factors 0.3, 0.3, 0.1, 0.1, 0.1 and 0.1 (genome fraction, largest alignment, NGA50, duplication ratio, number of contigs, and number of mismatches per 100 kb), respectively. The score of an assembler with metric *i* was formulated based on the scale average performance *sp*_*i*_ and then multiplied by a factor of 10 to ensure the score is in the range of 0-10:

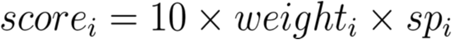

where *sp*_*i*_ is the scaled performance for metric *i*. The value was scaled into 0-1 with min-max normalization defined as follows:

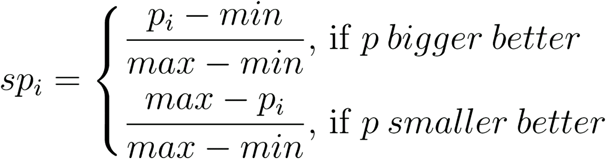

In the formula, *p*_*i*_ is the average performance across all samples of the given assembler for metric *i* and the min and max are the smallest and largest average performance value on metric *i* among all assemblers.

### Determination of genome differences between two strains

MUMmer v3.23 with default setting was used to align two genomes of the strains in each mixture and to identify the differences between genomes as ground truth. Command “show-snps” of the MUMmer package was employed to determine the SNPs and short InDels differing between two aligned genomes with parameter setting “--CTHr”, where the repeat regions were masked. The genomic differences between TB40/E and Merlin were considered as the ground truth variants for the TM mixtures, while differences between TB40/E and AD169 were considered as the ground truth for the TA mixtures.

### Variant calling

Quality controlled reads were mapped against the reference genome of the HCMV strains Merlin and AD169 using BWA-MEM with a seed length of 31. HCMV Merlin and AD169 genomes were used as reference genomes, as they were the major strains in all mixtures. The resulting BAM files were deduplicated with the Picard package (http://broadinstitute.github.io/picard/) to remove possible amplification duplicates that may bias the allele frequency of identified variants. To compare the performance of different variant callers, we used LoFreq (parameter: -q 20 -Q 20 -m 20), VarScan2 (---min-avg-qual 20 --p-value 0.01), FreeBayes (--p 1 -m 20 -q 20 -F 0.01 --min-coverage 10), CLC (overall read depth ≥10, average basecall quality ≥20, forward/reverse read balance 0.1-0.9 and variant frequency ≥0.1%), BCFtools (--p --ploidy 1 -mv -Ob) and GATK HaplotypeCaller (--min-base-quality-score 20 - ploidy 1) to identify variants. The variants from the difference between genomes detected by MUMmer were considered as positive variants. Based on this standard, precision, recall, and F1-score were computed to evaluate those callers. The pairwise genome differences of 30 *E. coli* or 30 HIV genomes were determined by MUMmer as well. To evaluate the performance of different callers for SNP and InDel prediction, the command vcfeval in RTG-tools [60] was used to generate recall-precision curves based on the Phred scaled “QUAL” score field (--squash-ploidy -f QUAL --sample ALT).

## Data and code availability

The benchmarking program developed in this study is available under the GNU General Public License V3.0 at https://github.com/hzi-bifo/Quasimodo. This program can be also used to assess variant calling and assembly results for other viral mixed strain data sets (see readme of the repository for details). All assembly and variant calling results are freely accessible on Zenodo (10.5281/zenodo.3739874). The sequence data were deposited in ENA with accession number PRJEB32127.

## Discussion and conclusions

Mixed infections with multiple HCMV strains are commonly observed in patients with active HCMV replication [10,17–20]. Accurately reconstructing the genomic sequences of the individual haplotypes has implications for gaining a deeper understanding of viral pathogenicity and viral diversity within the host. To identify the most suitable software for analysis of mixed viral genome sequencing samples with low evolutionary divergence and comparatively large genomes, we evaluated multiple state-of-the-art assemblers and variant callers on lab-generated strain mixtures of HCMV.

In the assembly benchmarking, most metagenome and genome assemblers, in particular metaSPAdes and IVA, recovered the abundant strains well in terms of metrics such as genome fraction, contig length and mismatches. When also considering strains of low abundance, Savage recovered the largest fractions of both underlying genomes in the reference-based mode. However, this was achieved in a highly fragmented manner, consistent with reports by the authors (Table 3 [34]). Thus, the state-of-the-art in assembly methods, including both generic (meta-)genome and specialized viral quasispecies assemblers, does not yet reconstruct large viral HCMV genomes of low abundance and low variant density with high quality. This may not be surprising since these programs were originally designed primarily for mixtures of large and much more divergent microbial genomes, or for viral genomes with a tenth of the size of the HCMV genome, but a higher variant density. In terms of resource usage, Ray and ABySS produced the smallest outputs, while megahit was the most memory efficient, as well as fastest assembler with good performance (weighted score >5).

Of the variant callers, LoFreq most faithfully identified only true variants across all samples, closely followed by VarScan2. Both had high F1-scores even on the samples with high mixing ratios. When analyzing the genomic context of the predicted variants, for true positive calls, we observed a context independent enrichment of T to C and C to T transitions. A preference for transitions over transversions is common in molecular evolution [61,62]. This is the case in terms of observed mutations and because transitions more often lead to synonymous mutations that tend to be neutral, rather than under negative selection, as most nonsynonymous changes on the population level.

For false variant calls, we found a striking enrichment of T to G changes in a G.G context, representing an unreported context-dependent signal. Calls with this pattern had lower allele frequencies than true positive variant calls and were more pronounced in sample with more PCR cycles used (protocol 2, 6-14 cycles versus 4 in protocol 1), indicating a link to DNA amplification. Amplification error introduced in PCR cycles will accumulate exponentially and occur at frequencies that depend on when they were introduced: PCR-induced errors are mostly of lower frequency unless introduced in one of the very early amplification cycles [57]. Schirmer and co-workers studied the error profiles for the amplicon sequencing using MiSeq with different library preparation methods and showed that the library preparation method and the choice of primers are the most significant sources of bias and cause distinct error patterns [63]. They also observed a run-specific preference for the substituting nucleotide. They observed that A and C were more prone to substitution errors (A to C and C to A) compared to G and T, which differ from our results. We could not find the context dependent signal for an HIV quasispecies data set that had been generated with Nextera XT DNA Library Prep chemistry (Illumina) on Illumina’s MiSeq platform, suggesting that the false positive pattern originates from a step unique to the HCMV sequencing protocol, such as pre-amplification and amplification PCR during library preparation.

Notably, the experimental protocols substantially affected the nature of the generated data and bioinformatics results. Protocol 1 led to substantial amplification of *E. coli* host DNA and thus lower coverage of the viral strains. This, together with the resulting differences in actual mixing ratios relative to protocol 2 likely explain the higher recall and slightly lower precision observed in variant detection (Figure S4). An earlier study based on simulated sequencing data also showed that variant calling on lower coverage samples achieved higher recall and lower precision [64]. Protocol 2 used more extensive DNA amplification together with cultivation in human cell culture. This resulted in higher coverage of viral strain genomes in comparison to protocol 1, and the doubling of context-dependent false positive variant calls within a G.G context discussed above (Figure 7).

Taken together, our results suggest that for strain mixtures of large DNA viruses with low variant density, many assemblers reconstruct the abundant strain with high quality, but assembly of the low abundant strains is still challenging. Variant callers designed for low frequency variant detection provided the best results and detected most true variants. These findings are relevant for the interpretation of program outputs when analyzing clinical patient samples. We also provide a resource that facilitates further benchmarking, including our result evaluation and visualization software QuasiModo, all produced benchmarking data sets and results, for flexible assessment of further methods on these and similar data sets.

## Key points

- The strain-resolved *de novo* assembly of large DNA virus with low variant density is challenging to all evaluated assemblers. Some generic (meta-)genome assemblers, such as metaSPAdes and IVA, performed particularly well in recovering the dominant strain.
- LoFreq and VarScan2 are good choices for identifying low frequency variants from strain mixture of large DNA viruses.
- The pattern of false variant calls likely links to the experimental protocol used to generate the sequencing data. More amplification cycles led to more pronounced false positive variant calls.
- All the analyses can be reproduced using QuasiModo developed in this study. QuasiModo can be also utilized to evaluate other methods using the benchmarking data sets in this study or similar data sets.

## Acknowledgments

We thank Dr. Till-Robin Lesker and Susanne Reimering for helpful comments. This work is supported by German Center for Infection Research (DZIF), Hannover-Braunschweig site, TI Bioinformatics Platform and TTU Infections of the Immunocompromised Host, as well as by the Deutsche Forschungsgemeinschaft through the Collaborative Research Center 900 “Chronic Infections”. A. Dhingra and J. Götting were supported by the graduate program “Infection Biology” of the Hannover Biomedical Research School.

## Description of the authors

**Z.-L. Deng** is a post-doctoral researcher at the Department Computational Biology of Infection Research of the Helmholtz Centre for Infection Research. His research focuses on viral genomics and microbiome.

**A. Dhingra** was a PhD student at the Institute of Virology in Hannover Medical School working on next generation sequencing of viruses. He is currently a post-doctoral researcher.

**A. Fritz** is a PhD student at the Department Computational Biology of Infection Research of the Helmholtz Centre for Infection Research. He focuses on developing software for strain-resolved assembly and metagenomics.

**J. Götting** is a PhD student at the Institute of Virology in Hannover Medical School working on next generation sequencing of viruses and NGS data analysis.

**P. C. Münch** is a PhD student at the Department Computational Biology of Infection Research of the Helmholtz Centre for Infection Research and Max von Pettenkofer Institute in Ludwig Maximilian University of Munich.

**L. Steinbrück** is working on next generation sequencing at the Institute of Virology in Hannover Medical School.

**T. F. Schulz** is the director of Institute of Virology in Hannover Medical School. And his research focuses on virology and infection.

**T. Ganzenmüller** was working as a clinical virologist at the Institute of Virology in Hannover Medical School. She is currently working at the Institute for Medical Virology in University Hospital Tübingen as a senior physician.

**A. C. McHardy** is the head of Department Computational Biology of Infection Research of the Helmholtz Centre for Infection Research. The department studies the human microbiome, viral and bacterial pathogens, and human cell lineages within individual patients by analysis of large-scale biological and epidemiological data sets with computational techniques.

## Notes

https://github.com/hzi-bifo/Quasimodo

### Competing Interest Statement

The authors have declared no competing interest.

