## Supplementary for "Evaluating assembly and variant calling software for strain-resolved analysis of large DNA-viruses"

#### Supplementary Material

##### Supplementary figures

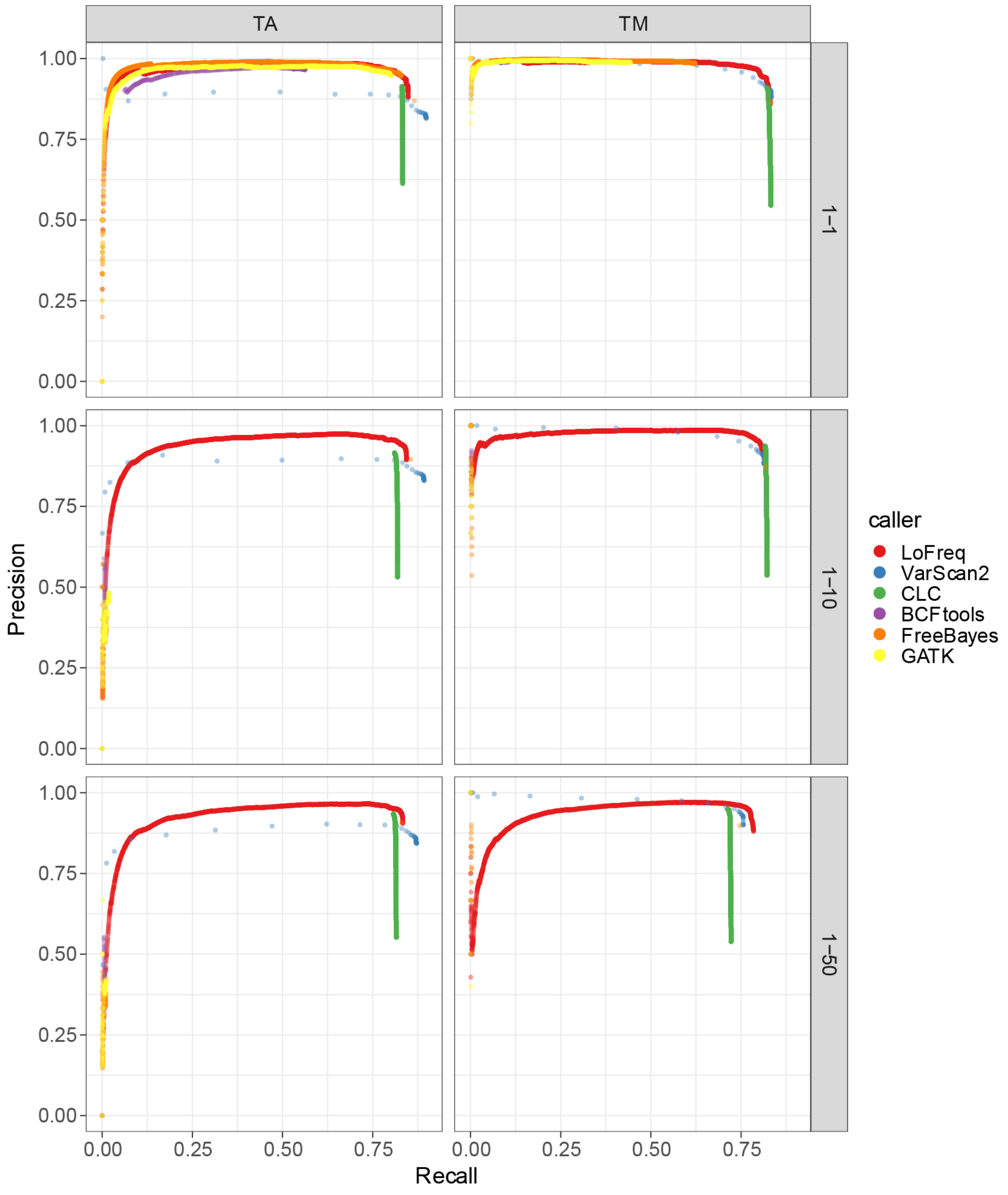


**Figure S1.** Comparison of variant callers using recall-precison curves across all reported scores for each variant caller across all variants (SNPs and INDELs).

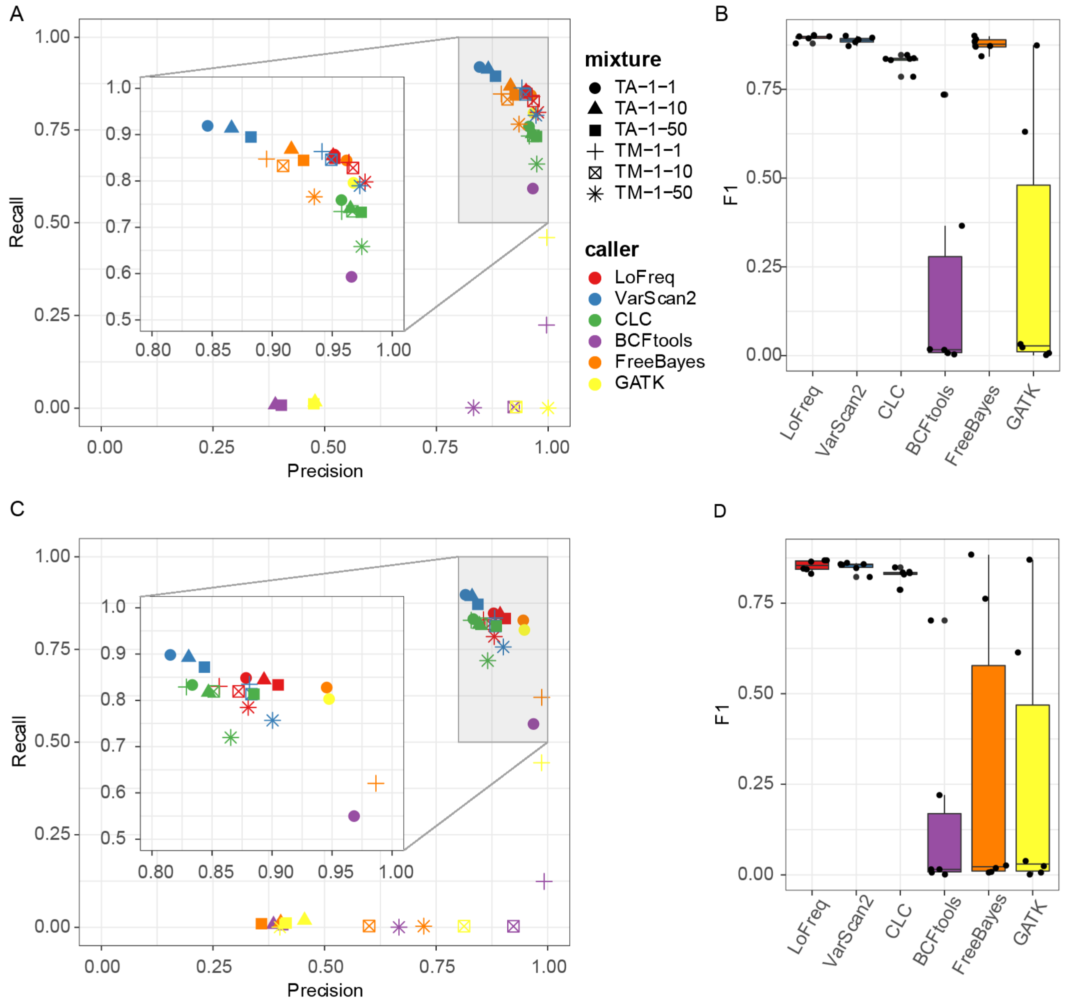


**Figure S2. Comparison of variants caller's performances.** (A) and (B) SNP calling performance with best F1-score. QUAL score thresholds were chosen to maximize the F1-score for each sample and each caller. (C) and (D) Variant calling performance based on identified SNPs and InDels with QUAL score >=20.

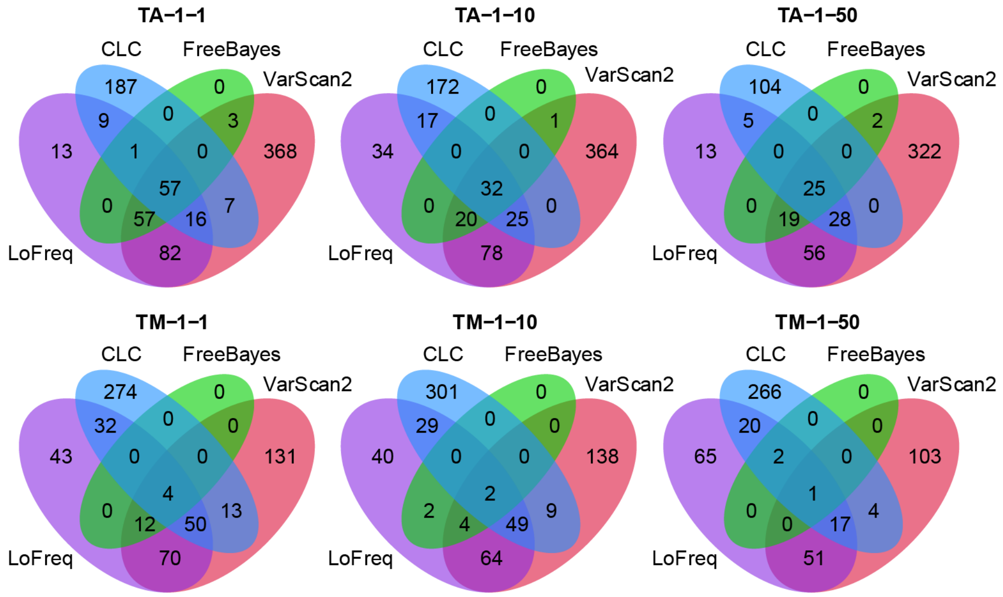


**Figure S3.** Comparison of FP SNPs identified by LoFreq, CLC, FreeBayes and VarScan2. The majority of FP SNPs predicted by LoFreq and FreeBayes were also identified by VarScan2.

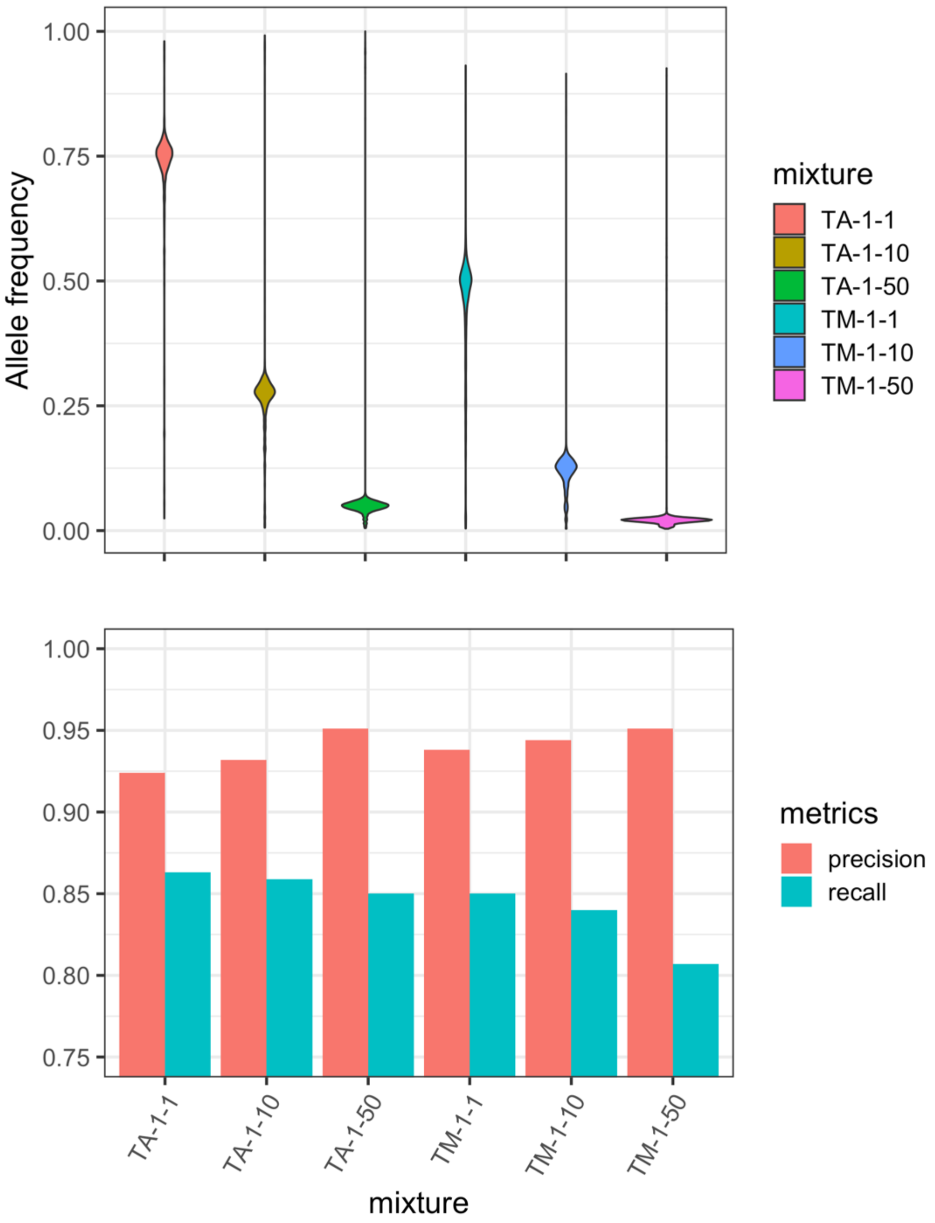


**Figure S4**. Allele frequency distribution of the true positive SNPs called by LoFreq strain mixtures and the precision and recall of LoFreq variant calls.

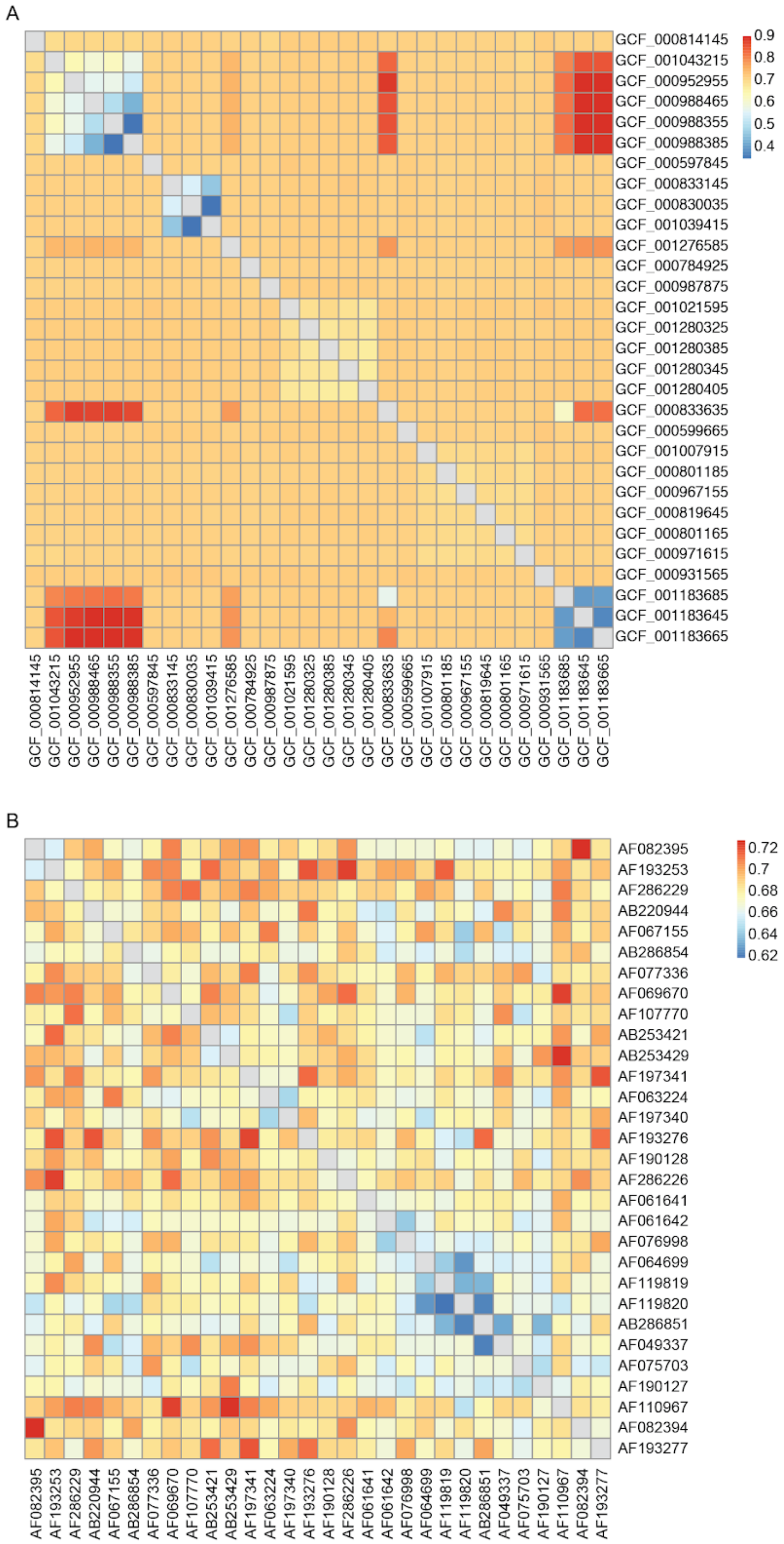


**Figure S5.** The fraction of transitions in all mutations between each pair of 30 *E. coli* genomes (A) and HIV genomes (B).

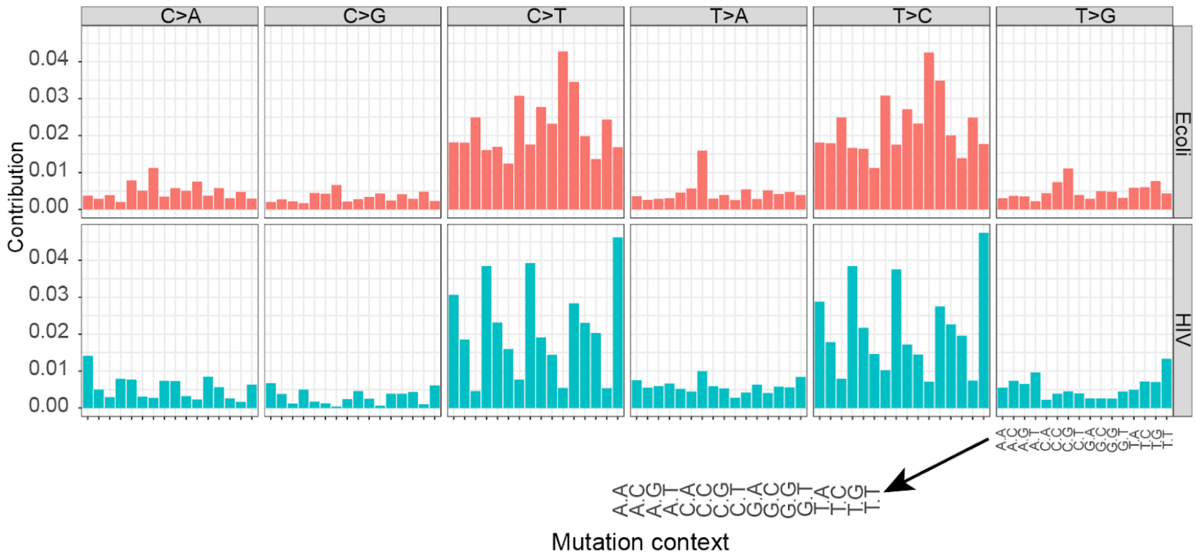


**Figure S6.** Average mutation contribution in different genomic context of variants for all pairs of 30 *E. coli* (upper panel) and 30 HIV genomes (bottom panel). The 30 *E. coli* and 30 HIV genomes were downloaded from NCBI refseq database.

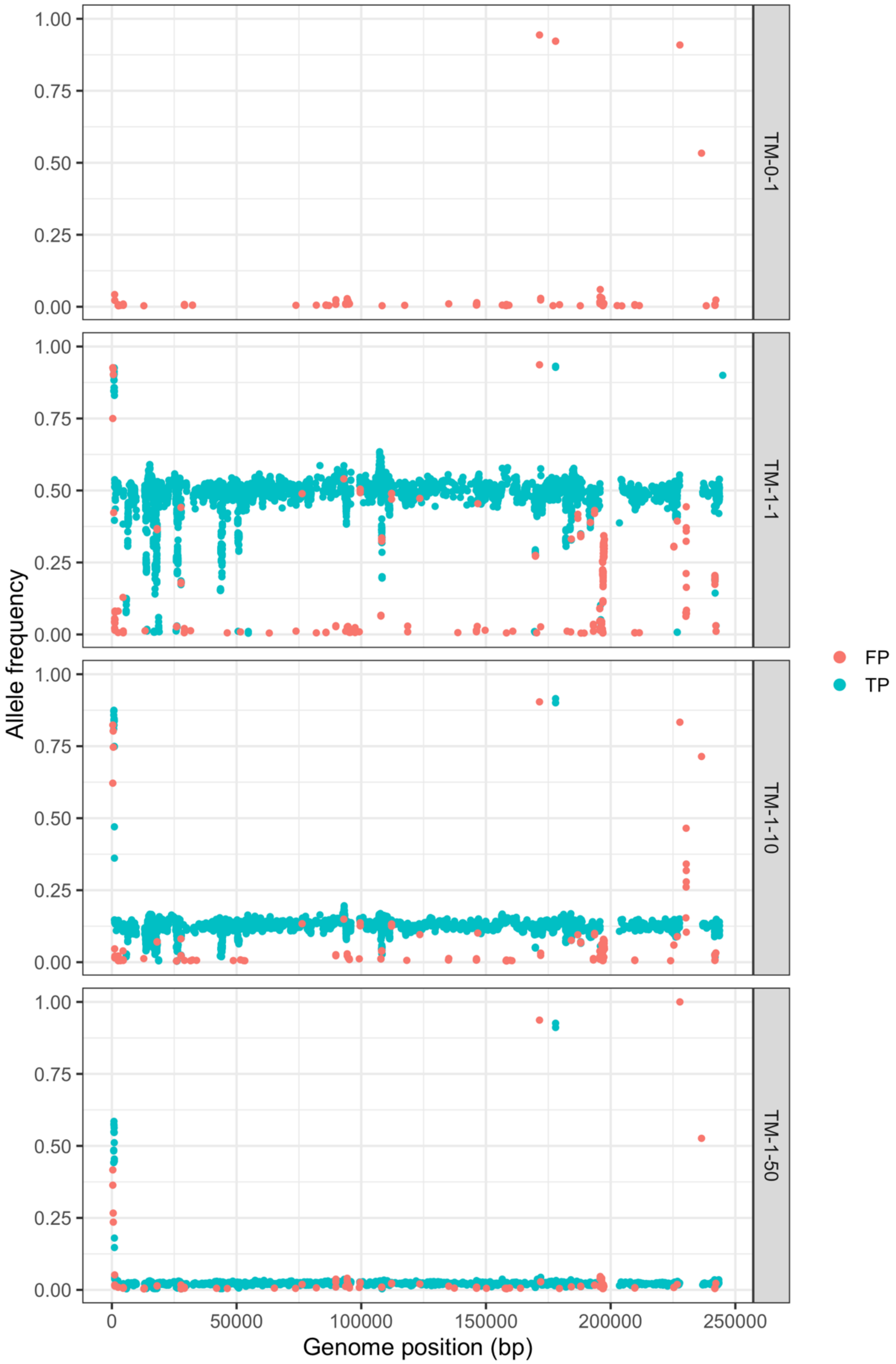


**Figure S7.** The allele frequency and distribution of SNPs identified by LoFreq on the sample from mixture TM and pure Merlin.

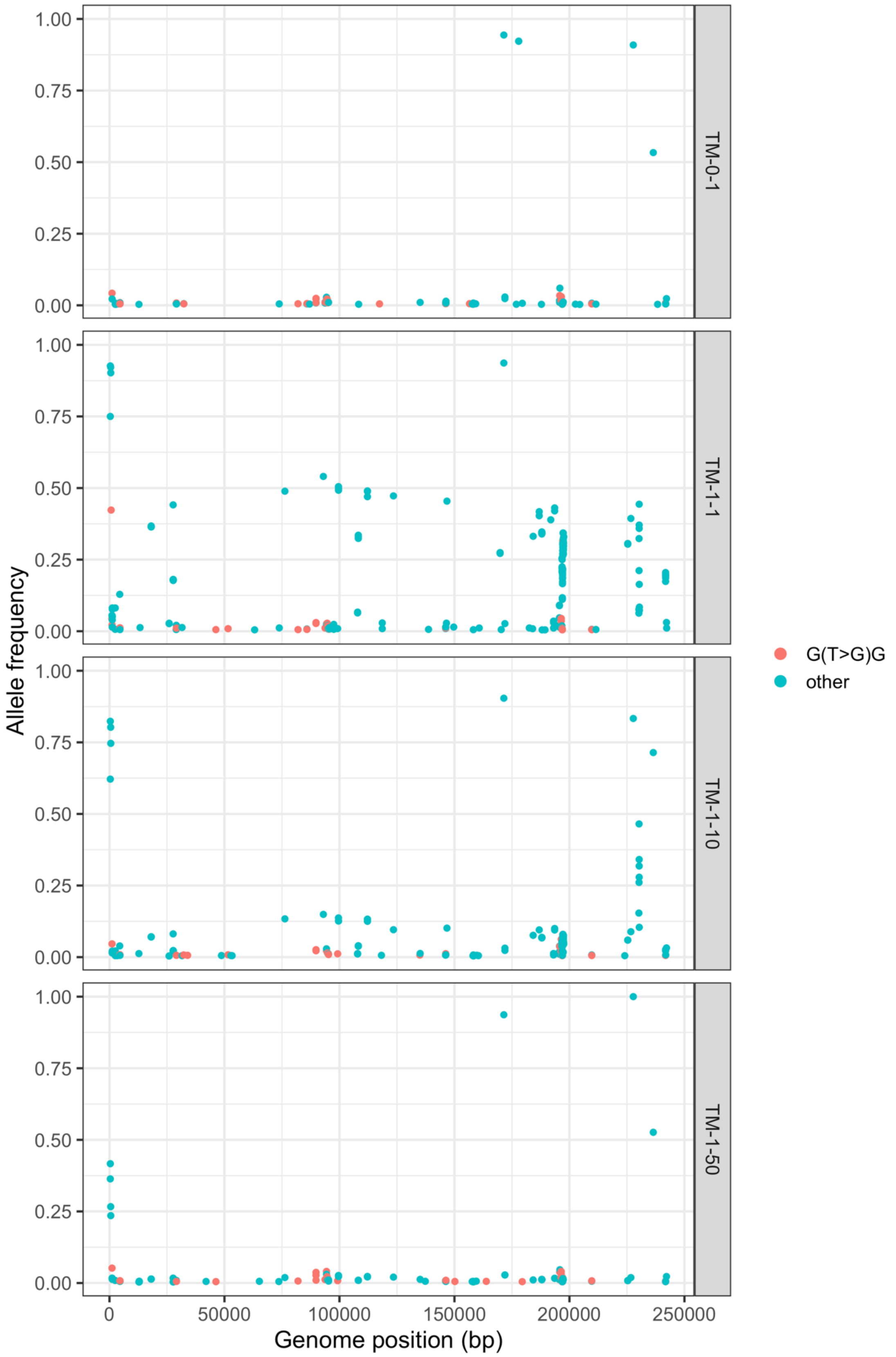


**Figure S8.** The allele frequency and distribution of FP SNPs identified by LoFreq on the sample from mixture TM and pure Merlin. T to G substitutions in the context of G.G are shown in red.

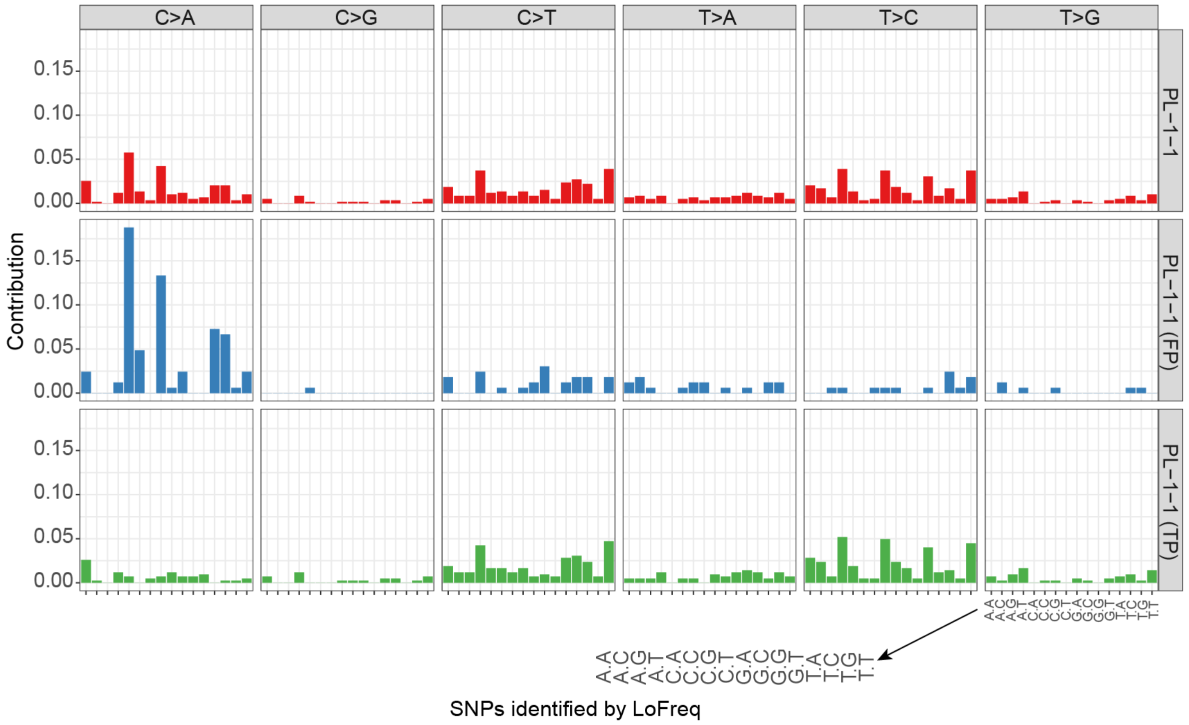


**Figure S9.** Genomic context pattern for the HIV PL-1-1 mixture.

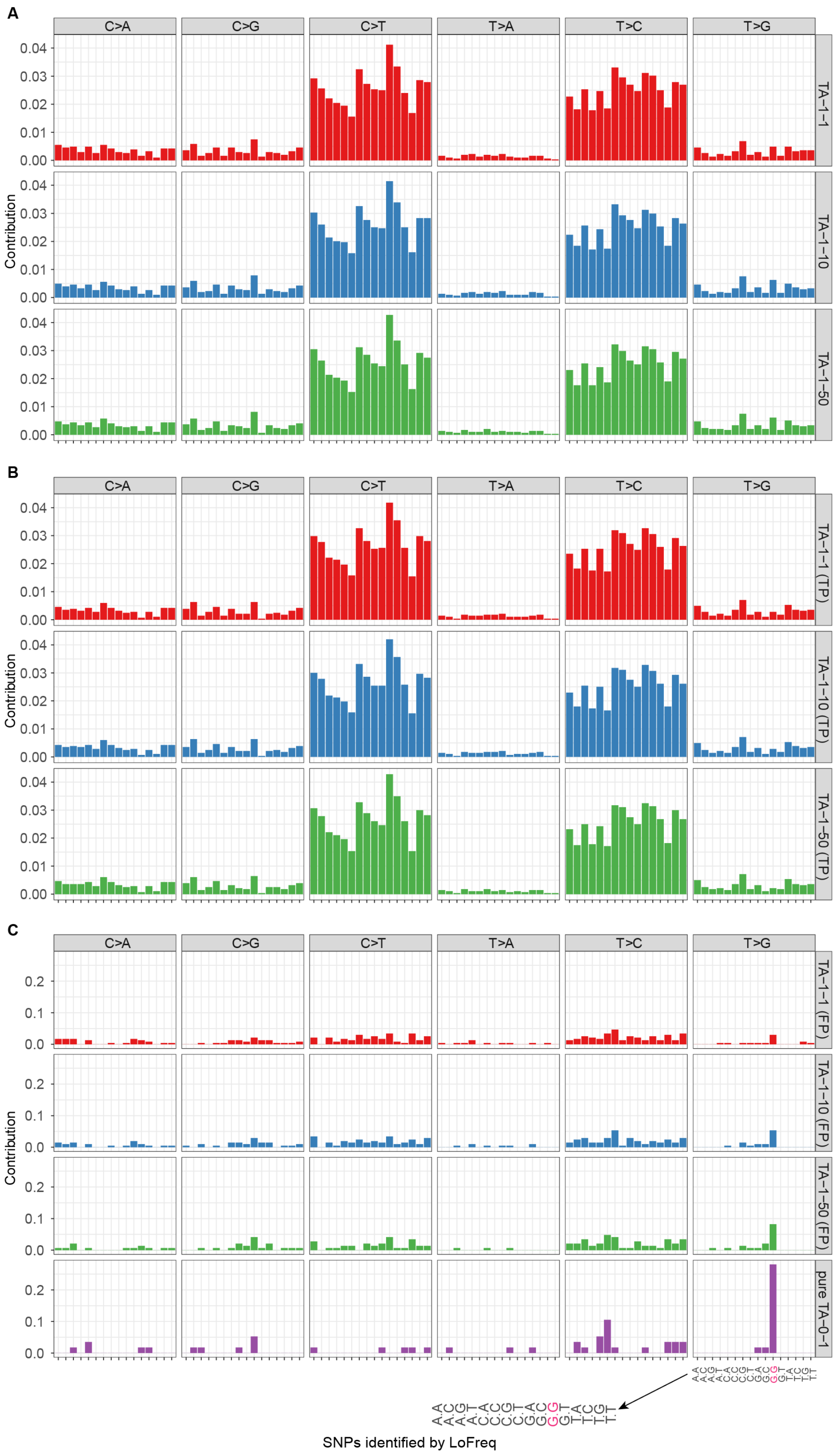


**Figure S10.** The mutation context of all identified SNPs (upper panel), true positive SNPs, and false positive SNPs for TA mixture. The SNPs were called by LoFreq.

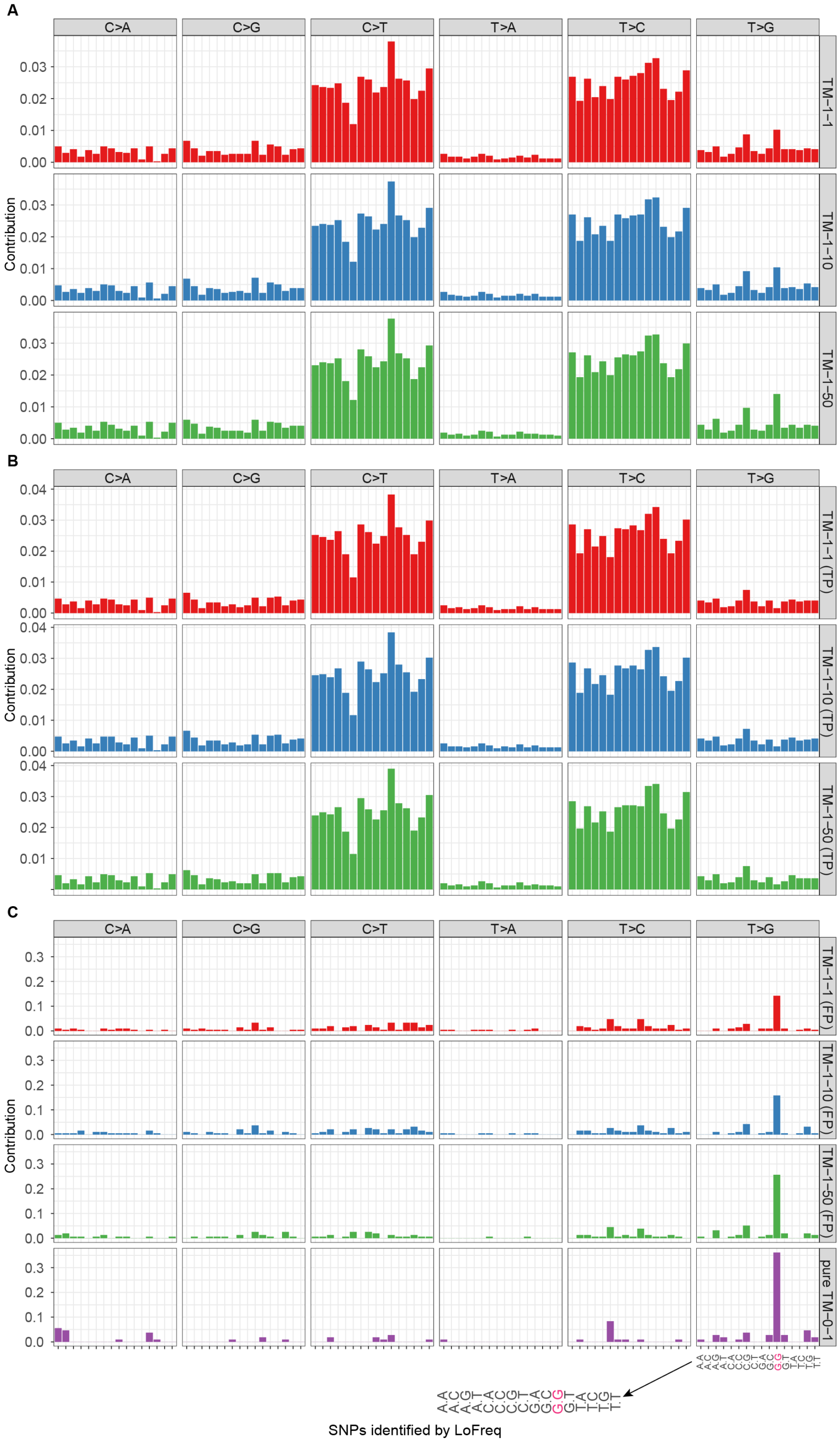


**Figure S11.** The mutation context of all identified SNPs (upper panel), true positive SNPs, and false positive SNPs for TM mixture. The SNPs were called by LoFreq.

### Supplementary tables

**Table S1** ANI between pairs of HCMV genomes.

| | **Merlin** | **AD169** | **TB40** |
|---|---|---|---|
| **Merlin** | 1 | 0.978 | 0.978 |
| **AD169** | 0.977 | 1 | 0.976 |
| **TB40** | 0.977 | 0.979 | 1 |

**Table S2**. Sequence sample sizes and read mapping rates.

| **Sample** | **Number reads for mapping** | ***E. coli* reads** | **Human reads** | **HCMV ref** | **HCMV mapped reads** |
|---|---|---|---|---|---|
| TA-0-1 | 1,927,695 | 2,372 | 483,998 | AD169 | 74.94% |
| TA-1-0 | 1,746,264 | 2,098 | 928,083 | TB40E | 46.70% |
| TA-1-10 | 1,709,083 | 1,407 | 937,308 | AD169 | 43.73% |
| TA-1-1 | 1,595,119 | 1,200 | 1,100,035 | AD169 | 28.56% |
| TA-1-50 | 1,475,223 | 2,011 | 603,575 | AD169 | 58.55% |
| TM-0-1 | 2,033,786 | 18,142 | 55 | Merlin | 99.32% |
| TM-1-0 | 842,943 | 435,400 | 2,456 | TB40E | 45.27% |
| TM-1-10 | 1,401,688 | 8,230 | 560 | Merlin | 98.75% |
| TM-1-1 | 1,379,143 | 9,609 | 1,688 | Merlin | 97.34% |
| TM-1-50 | 1,645,708 | 14,736 | 95 | Merlin | 99.18% |

**Table S3**. Assembly quality according to different MetaQUAST metrics

| **Assembler** | **Metrics** | **Average** | **SD** | **Rank** |
|---|---|---|---|---|
| ABySS | # contigs | 14.8500 | 10.3415 | 4 |
| ABySS | # mismatches per 100 kbp | 152.7590 | 140.9024 | 9 |
| ABySS | Duplication ratio | 1.0304 | 0.0237 | 5 |
| ABySS | Genome fraction (%) | 58.2991 | 9.2640 | 4 |
| ABySS | Largest alignment | 90829.0000 | 61619.4993 | 5 |
| ABySS | NGA50 | 69858.9500 | 78361.4076 | 4 |
| IDBA | # contigs | 98.0500 | 33.1675 | 8 |
| IDBA | # mismatches per 100 kbp | 88.4130 | 63.0756 | 7 |
| IDBA | Duplication ratio | 1.0306 | 0.0119 | 6 |
| IDBA | Genome fraction (%) | 33.2301 | 10.0549 | 10 |
| IDBA | Largest alignment | 8360.6000 | 5756.1502 | 10 |

| | | | | |
|---|---|---|---|---|
| IDBA | NGA50 | 807.7000 | 1528.1691 | 10 |
| IVA | # contigs | 8.0500 | 7.9213 | 1 |
| IVA | # mismatches per 100 kbp | 74.9910 | 151.1568 | 6 |
| IVA | Duplication ratio | 1.0122 | 0.0261 | 3 |
| IVA | Genome fraction (%) | 45.2945 | 12.3756 | 8 |
| IVA | Largest alignment | 159583.8000 | 77840.4515 | 1 |
| IVA | NGA50 | 118449.8000 | 80726.2336 | 1 |
| megahit | # contigs | 274.1000 | 229.1707 | 10 |
| megahit | # mismatches per 100 kbp | 1709.2940 | 1683.7245 | 10 |
| megahit | Duplication ratio | 1.7289 | 0.6251 | 10 |
| megahit | Genome fraction (%) | 60.6160 | 11.3693 | 2 |
| megahit | Largest alignment | 111325.8000 | 55683.7843 | 3 |
| megahit | NGA50 | 72387.0500 | 68605.9038 | 3 |
| metaSPAdes | # contigs | 12.4500 | 8.9859 | 2 |
| metaSPAdes | # mismatches per 100 kbp | 114.7410 | 266.9891 | 8 |
| metaSPAdes | Duplication ratio | 1.0073 | 0.0096 | 2 |
| metaSPAdes | Genome fraction (%) | 54.5381 | 6.3756 | 5 |
| metaSPAdes | Largest alignment | 145854.3000 | 56616.2594 | 2 |
| metaSPAdes | NGA50 | 102292.6500 | 68972.8802 | 2 |
| Ray | # contigs | 12.5000 | 10.9671 | 3 |
| Ray | # mismatches per 100 kbp | 43.4630 | 56.8234 | 4 |
| Ray | Duplication ratio | 1.0527 | 0.0496 | 8 |
| Ray | Genome fraction (%) | 49.8487 | 2.7212 | 6 |
| Ray | Largest alignment | 92791.4000 | 67899.5739 | 4 |
| Ray | NGA50 | 62380.8500 | 66970.7912 | 5 |
| Savage | # contigs | 200.4000 | 133.2007 | 9 |
| Savage | # mismatches per 100 kbp | 39.4990 | 42.7163 | 2 |
| Savage | Duplication ratio | 1.3766 | 0.2297 | 9 |
| Savage | Genome fraction (%) | 64.3606 | 27.1913 | 1 |
| Savage | Largest alignment | 21502.5000 | 22764.8772 | 8 |
| Savage | NGA50 | 6820.1000 | 11633.4454 | 8 |
| SPAdes | # contigs | 55.7000 | 40.6695 | 7 |
| SPAdes | # mismatches per 100 kbp | 39.5100 | 41.5310 | 3 |
| SPAdes | Duplication ratio | 1.0341 | 0.0267 | 7 |
| SPAdes | Genome fraction (%) | 58.5633 | 9.4669 | 3 |
| SPAdes | Largest alignment | 72552.6000 | 78224.1194 | 6 |
| SPAdes | NGA50 | 58068.1500 | 87597.4403 | 6 |
| tadpole | # contigs | 34.0500 | 13.6044 | 5 |
| tadpole | # mismatches per 100 kbp | 32.2460 | 54.3644 | 1 |
| tadpole | Duplication ratio | 1.0014 | 0.0011 | 1 |
| tadpole | Genome fraction (%) | 33.7846 | 15.6001 | 9 |
| tadpole | Largest alignment | 29431.7000 | 21598.7536 | 7 |

| | | | | |
|---|---|---|---|---|
| tadpole | NGA50 | 10890.2000 | 15426.9931 | 7 |
| Vicuna | # contigs | 36.7000 | 10.1440 | 6 |
| Vicuna | # mismatches per 100 kbp | 70.6150 | 69.0543 | 5 |
| Vicuna | Duplication ratio | 1.0186 | 0.0059 | 4 |
| Vicuna | Genome fraction (%) | 47.2257 | 0.9994 | 7 |
| Vicuna | Largest alignment | 20307.4000 | 8070.6917 | 9 |
| Vicuna | NGA50 | 6522.6000 | 5790.0728 | 9 |

**Table S4**. Summarized scores for the assemblers

| Assembler | Weighted score |
|---|---|
| ABySS | 7.502 |
| IDBA | 2.588 |
| IVA | 8.122 |
| megahit | 5.290 |
| metaSPAdes | 8.570 |
| Ray | 6.706 |
| Savage | 5.069 |
| SPAdes | 6.973 |
| tadpole | 3.459 |
| Vicuna | 4.480 |

**Table S5.** Performance metrics for variants callers LoFreq, VarScan2, FreeBayes, BCFtools, CLC and GATK on all samples

| Caller | Sample | Genome | Call | TP | FP | Precision | Recall | F1 |
|---|---|---|---|---|---|---|---|---|
| BCFtools | TA-0-1 | 0 | 25 | 0 | 25 | 0 | NA | NA |
| BCFtools | TA-1-0 | 0 | 1 | 0 | 1 | 0 | NA | NA |
| BCFtools | TA-1-10 | 3301 | 70 | 27 | 43 | 0.386 | 0.008 | 0.016 |
| BCFtools | TA-1-1 | 3301 | 1978 | 1916 | 62 | 0.969 | 0.58 | 0.726 |
| BCFtools | TA-1-50 | 3301 | 64 | 26 | 38 | 0.406 | 0.008 | 0.016 |
| BCFtools | TM-0-1 | 0 | 3 | 0 | 3 | 0 | NA | NA |
| BCFtools | TM-1-0 | 0 | 2 | 0 | 2 | 0 | NA | NA |
| BCFtools | TM-1-10 | 3784 | 13 | 12 | 1 | 0.923 | 0.003 | 0.006 |
| BCFtools | TM-1-1 | 3784 | 501 | 498 | 3 | 0.994 | 0.132 | 0.233 |
| BCFtools | TM-1-50 | 3784 | 3 | 2 | 1 | 0.667 | 0.001 | 0.002 |

| | | | | | | | | |
|---|---|---|---|---|---|---|---|---|
| CLC | TA-0-1 | 0 | 102 | 0 | 102 | 0 | NA | NA |
| CLC | TA-1-0 | 0 | 94 | 0 | 94 | 0 | NA | NA |
| CLC | TA-1-10 | 3301 | 2703 | 2457 | 246 | 0.909 | 0.744 | 0.818 |
| CLC | TA-1-1 | 3301 | 2792 | 2515 | 277 | 0.901 | 0.762 | 0.826 |
| CLC | TA-1-50 | 3301 | 2593 | 2431 | 162 | 0.938 | 0.736 | 0.825 |
| CLC | TM-0-1 | 0 | 359 | 0 | 359 | 0 | NA | NA |
| CLC | TM-1-0 | 0 | 175 | 0 | 175 | 0 | NA | NA |
| CLC | TM-1-10 | 3784 | 3177 | 2787 | 390 | 0.877 | 0.737 | 0.801 |
| CLC | TM-1-1 | 3784 | 3170 | 2797 | 373 | 0.882 | 0.739 | 0.804 |
| CLC | TM-1-50 | 3784 | 2819 | 2509 | 310 | 0.89 | 0.663 | 0.76 |
| FreeBayes | TA-0-1 | 0 | 30 | 0 | 30 | 0 | NA | NA |
| FreeBayes | TA-1-0 | 0 | 1 | 0 | 1 | 0 | NA | NA |
| FreeBayes | TA-1-10 | 3301 | 96 | 43 | 53 | 0.448 | 0.013 | 0.025 |
| FreeBayes | TA-1-1 | 3301 | 2906 | 2788 | 118 | 0.959 | 0.845 | 0.898 |
| FreeBayes | TA-1-50 | 3301 | 77 | 31 | 46 | 0.403 | 0.009 | 0.018 |
| FreeBayes | TM-0-1 | 0 | 5 | 0 | 5 | 0 | NA | NA |
| FreeBayes | TM-1-0 | 0 | 2 | 0 | 2 | 0 | NA | NA |
| FreeBayes | TM-1-10 | 3784 | 23 | 15 | 8 | 0.652 | 0.004 | 0.008 |
| FreeBayes | TM-1-1 | 3784 | 2451 | 2435 | 16 | 0.993 | 0.643 | 0.781 |
| FreeBayes | TM-1-50 | 3784 | 16 | 13 | 3 | 0.812 | 0.003 | 0.006 |
| GATK | TA-0-1 | 0 | 36 | 0 | 36 | 0 | NA | NA |
| GATK | TA-1-0 | 0 | 1 | 0 | 1 | 0 | NA | NA |
| GATK | TA-1-10 | 3301 | 116 | 55 | 61 | 0.474 | 0.017 | 0.033 |
| GATK | TA-1-1 | 3301 | 2719 | 2631 | 88 | 0.968 | 0.797 | 0.874 |
| GATK | TA-1-50 | 3301 | 84 | 39 | 45 | 0.464 | 0.012 | 0.023 |
| GATK | TM-0-1 | 0 | 4 | 0 | 4 | 0 | NA | NA |
| GATK | TM-1-0 | 0 | 2 | 0 | 2 | 0 | NA | NA |
| GATK | TM-1-10 | 3784 | 14 | 13 | 1 | 0.929 | 0.003 | 0.006 |
| GATK | TM-1-1 | 3784 | 1746 | 1741 | 5 | 0.997 | 0.46 | 0.63 |
| GATK | TM-1-50 | 3784 | 4 | 2 | 2 | 0.5 | 0.001 | 0.002 |
| LoFreq | TA-0-1 | 0 | 57 | 0 | 57 | 0 | NA | NA |
| LoFreq | TA-1-0 | 0 | 48 | 0 | 48 | 0 | NA | NA |
| LoFreq | TA-1-10 | 3301 | 3040 | 2834 | 206 | 0.932 | 0.859 | 0.894 |
| LoFreq | TA-1-1 | 3301 | 3084 | 2849 | 235 | 0.924 | 0.863 | 0.892 |
| LoFreq | TA-1-50 | 3301 | 2951 | 2805 | 146 | 0.951 | 0.85 | 0.898 |

| | | | | | | | | |
|---|---|---|---|---|---|---|---|---|
| LoFreq | TM-0-1 | 0 | 108 | 0 | 108 | 0 | NA | NA |
| LoFreq | TM-1-0 | 0 | 30 | 0 | 30 | 0 | NA | NA |
| LoFreq | TM-1-10 | 3784 | 3370 | 3180 | 190 | 0.944 | 0.84 | 0.889 |
| LoFreq | TM-1-1 | 3784 | 3427 | 3216 | 211 | 0.938 | 0.85 | 0.892 |
| LoFreq | TM-1-50 | 3784 | 3210 | 3054 | 156 | 0.951 | 0.807 | 0.873 |
| VarScan2 | TA-0-1 | 0 | 114 | 0 | 114 | 0 | NA | NA |
| VarScan2 | TA-1-0 | 0 | 47 | 0 | 47 | 0 | NA | NA |
| VarScan2 | TA-1-10 | 3301 | 3545 | 3025 | 520 | 0.853 | 0.916 | 0.883 |
| VarScan2 | TA-1-1 | 3301 | 3631 | 3041 | 590 | 0.838 | 0.921 | 0.878 |
| VarScan2 | TA-1-50 | 3301 | 3411 | 2959 | 452 | 0.867 | 0.896 | 0.881 |
| VarScan2 | TM-0-1 | 0 | 98 | 0 | 98 | 0 | NA | NA |
| VarScan2 | TM-1-0 | 0 | 26 | 0 | 26 | 0 | NA | NA |
| VarScan2 | TM-1-10 | 3784 | 3470 | 3202 | 266 | 0.923 | 0.846 | 0.883 |
| VarScan2 | TM-1-1 | 3784 | 3555 | 3274 | 280 | 0.921 | 0.865 | 0.892 |
| VarScan2 | TM-1-50 | 3784 | 3168 | 2991 | 176 | 0.944 | 0.79 | 0.86 |

**Table S6.** The QUAL score thresholds maximizing F1-score for each variant caller on each sample

| Caller | Mixture | Threshold | TP | FP | FN | Precision | Recall | F1 |
|---|---|---|---|---|---|---|---|---|
| LoFreq | TA-1-10 | 162 | 2908 | 203 | 559 | 0.9347 | 0.8396 | 0.8846 |
| VarScan2 | TA-1-10 | 39 | 3086 | 548 | 397 | 0.8492 | 0.8862 | 0.8673 |
| CLC | TA-1-10 | 200 | 2656 | 241 | 660 | 0.9168 | 0.8108 | 0.8606 |
| BCFtools | TA-1-10 | 6.046 | 30 | 47 | 3456 | 0.3896 | 0.0086 | 0.0168 |
| FreeBayes | TA-1-10 | 0 | 2945 | 345 | 502 | 0.8951 | 0.856 | 0.8751 |
| GATK | TA-1-10 | 54.01 | 66 | 78 | 3419 | 0.4583 | 0.0192 | 0.0369 |
| LoFreq | TA-1-1 | 241 | 2919 | 200 | 548 | 0.9359 | 0.8428 | 0.8869 |
| VarScan2 | TA-1-1 | 37 | 3113 | 642 | 364 | 0.829 | 0.8956 | 0.861 |
| CLC | TA-1-1 | 200 | 2718 | 255 | 588 | 0.9142 | 0.8313 | 0.8708 |
| BCFtools | TA-1-1 | 5.292 | 1960 | 70 | 1526 | 0.9655 | 0.5622 | 0.7107 |
| FreeBayes | TA-1-1 | 915.155 | 2846 | 136 | 613 | 0.9544 | 0.8242 | 0.8845 |
| GATK | TA-1-1 | 35.01 | 2758 | 153 | 687 | 0.9474 | 0.8029 | 0.8692 |
| LoFreq | TA-1-50 | 168 | 2862 | 169 | 606 | 0.9442 | 0.8261 | 0.8812 |
| VarScan2 | TA-1-50 | 40 | 3007 | 464 | 476 | 0.8663 | 0.8636 | 0.865 |
| CLC | TA-1-50 | 200 | 2649 | 184 | 671 | 0.9351 | 0.8075 | 0.8666 |
| BCFtools | TA-1-50 | 9.702 | 27 | 40 | 3459 | 0.403 | 0.0077 | 0.0152 |
| FreeBayes | TA-1-50 | 0 | 2863 | 294 | 590 | 0.9069 | 0.8308 | 0.8671 |
| GATK | TA-1-50 | 85.01 | 41 | 56 | 3445 | 0.4227 | 0.0118 | 0.0229 |

| | | | | | | | | |
|---|---|---|---|---|---|---|---|---|
| LoFreq | TM-1-10 | 522 | 3213 | 171 | 801 | 0.9495 | 0.8011 | 0.869 |
| VarScan2 | TM-1-10 | 33 | 3270 | 326 | 758 | 0.9093 | 0.8117 | 0.8578 |
| CLC | TM-1-10 | 200 | 3061 | 204 | 741 | 0.9375 | 0.816 | 0.8725 |
| BCFtools | TM-1-10 | 26.626 | 12 | 1 | 4016 | 0.9231 | 0.003 | 0.0059 |
| FreeBayes | TM-1-10 | 0 | 3240 | 476 | 733 | 0.8719 | 0.818 | 0.8441 |
| GATK | TM-1-10 | 5975.01 | 13 | 2 | 4015 | 0.8667 | 0.0032 | 0.0064 |
| LoFreq | TM-1-1 | 459 | 3284 | 209 | 725 | 0.9402 | 0.8201 | 0.876 |
| VarScan2 | TM-1-1 | 34 | 3348 | 361 | 677 | 0.9027 | 0.8318 | 0.8658 |
| CLC | TM-1-1 | 200 | 3084 | 307 | 716 | 0.9095 | 0.8222 | 0.8637 |
| BCFtools | TM-1-1 | 4.247 | 849 | 4 | 3179 | 0.9953 | 0.2108 | 0.3479 |
| FreeBayes | TM-1-1 | 0 | 3297 | 513 | 672 | 0.8654 | 0.8332 | 0.849 |
| GATK | TM-1-1 | 35.01 | 1788 | 25 | 2239 | 0.9862 | 0.4441 | 0.6125 |
| LoFreq | TM-1-50 | 156 | 3109 | 186 | 907 | 0.9436 | 0.7749 | 0.851 |
| VarScan2 | TM-1-50 | 34 | 3043 | 234 | 985 | 0.9286 | 0.7555 | 0.8331 |
| CLC | TM-1-50 | 87.62 | 2715 | 153 | 1142 | 0.9467 | 0.7165 | 0.8156 |
| BCFtools | TM-1-50 | 4.042 | 5 | 1 | 4023 | 0.8333 | 0.0012 | 0.0025 |
| FreeBayes | TM-1-50 | 0 | 2974 | 337 | 1021 | 0.8982 | 0.7465 | 0.8154 |
| GATK | TM-1-50 | 122671.01 | 2 | 0 | 4026 | 1 | 5.00E-04 | 0.001 |

**Table S7.** The NCBI refseq accession numbers of *E. coli* and HIV genomes used to calculate the transition and transversion ratio

| ***E. coli*** **genomes** | **HIV genomes** |
|---|---|
| GCF_000597845 | AB220944 |
| GCF_000599665 | AB253421 |
| GCF_000784925 | AB253429 |
| GCF_000801165 | AB286851 |
| GCF_000801185 | AB286854 |
| GCF_000814145 | AF049337 |
| GCF_000819645 | AF061641 |
| GCF_000830035 | AF061642 |
| GCF_000833145 | AF063224 |
| GCF_000833635 | AF064699 |
| GCF_000931565 | AF067155 |
| GCF_000952955 | AF069670 |
| GCF_000967155 | AF075703 |
| GCF_000971615 | AF076998 |
| GCF_000987875 | AF077336 |
| GCF_000988355 | AF082394 |
| GCF_000988385 | AF082395 |
| GCF_000988465 | AF107770 |
| GCF_001007915 | AF110967 |

| | |
|---|---|
| GCF_001021595 | AF119819 |
| GCF_001039415 | AF119820 |
| GCF_001043215 | AF190127 |
| GCF_001183645 | AF190128 |
| GCF_001183665 | AF193253 |
| GCF_001183685 | AF193276 |
| GCF_001276585 | AF193277 |
| GCF_001280325 | AF197340 |
| GCF_001280345 | AF197341 |
| GCF_001280385 | AF286226 |
| GCF_001280405 | AF286229 |
